# *Streptococcus pyogenes* transforms the salivary proteome during controlled human infection

**DOI:** 10.64898/2026.10.20.760561

**Authors:** Anuk Dulanta Indraratna, Dezerae Cox, Plabon Kumar Das, Chi-Hung Lin, Kristy Azzopardi, Joshua Osowicki, Andrew Steer, Nicolle Packer, Danielle Skropeta, Arun Everest-Dass, Martina Sanderson-Smith

**Affiliations:** Molecular Horizons, School of Science, University of Wollongong, NSW; Institute for Biomedicine and Glycomics, Griffith University, QLD; Tropical Diseases, Murdoch Children’s Research Institute, VIC; Department of Paediatrics, University of Melbourne, VIC; Infectious Diseases Unit, The Royal Children’s Hospital Melbourne, VIC; MultiOmics Research Centre, School of Natural Sciences, Macquarie University, NSW

**Author notes:** These authors contributed equally (Arun Everest-Dass, Martina Sanderson-Smith). Corresponding authors &.

## Abstract

*Streptococcus pyogenes* (Group A *Streptococcus*, GAS) is among the leading causes of infectious mortality and primarily manifests as acute pharyngitis. The development of an effective vaccine continues to be hindered by an incomplete understanding of the host immune response, particularly in the oropharynx. Here, we provided detailed insights into the dynamic salivary proteome during acute illness in a human clinical trial of controlled infection of 20 healthy adults (CHIVAS-M75 trial). Deep profiling of salivary proteins via mass spectrometry revealed a striking increase in pro-inflammatory plasma proteins such as alpha1-antitrypsin and elements of the complement system during peak infection as a clear indication of the acute phase response in the oral environment. Many of these proteins, such as fibrinogen, are well-defined targets of key streptococcal virulence factors, with others, such as vitronectin, suggested to play a role in the host-pathogen relationship. Several salivary proteins with known antimicrobial properties, such as mucin-7 and cystatin, decreased universally across the infected cohort. For the first time, we have described the local and immediate protein response to acute pharyngitis in the oral cavity, highlighting a general host inflammatory signature and an abundance of GAS virulence targets infiltrating to the site of infection.

## Introduction

Group A Streptococcus (*Streptococcus pyogenes*, GAS) is a Gram-positive bacterial pathogen, responsible for a diverse range of human infections, spanning from superficial to life-threatening. GAS is the most common bacterial cause of pharyngitis, which is typically self-limiting and uncomplicated, but may also manifest as invasive, deeper tissue infections [1]. Post-infection auto-immune sequelae, notably rheumatic heart disease, contribute significantly to the global burden of disease. Prior to the emergence of SARS-CoV-2, GAS was considered among the top 10 communicable causes of human mortality [2].

Knowledge gaps regarding streptococcal pathogenesis have prevented a comprehensive understanding of the human response to GAS infection, impeding vaccine research. To this end, a ‘roadmap’ of priority activities to accelerate vaccine development has been proposed by the World Health Organisation [3], including the now-realised Controlled Human Infection for Vaccination Against Streptococcus (or CHIVAS-M75) model [4, 5]. Human infection models circumvent the inherent and GAS-specific limitations of animal models, enabling unparalleled interrogation of the differences between diseased and healthy samples, even in small cohorts and in highly variable tissue, such as saliva. Several, distinct biological insights have arisen from CHIVAS-M75 and related models (Table S1) [6–14], including identification of a consistent inflammatory signature in serum and a prolonged serum antibody response.

Although studies have focused primarily on circulating factors rather than the mucosal environment, targeted flow cytometric analysis of saliva did reveal an increase in IL-6 at 24 h from baseline in participants with pharyngitis [8]. However, interpretations of saliva data were generally constrained by high variability and low absolute signal, suggesting a limitation of this technique. Such challenges highlight the need for orthogonal molecular platforms capable of deeper, more sensitive quantitation such as mass spectrometry. Importantly, untargeted analysis for hypothesis-generation in the context of CHIVAS has not been performed to date, particularly with respect to the mucosal environment, providing an opportunity for novel investigation which may reveal unexpected biosignatures previously unidentified in GAS pathophysiology.

Saliva is an increasingly appreciated substrate in basic and clinical research contexts [15, 16]. Its collection is non-invasive and can be performed repeatedly and without medical training [17]. In the realm of infectious disease, saliva is an easily accessible diagnostic biofluid for oral infections [9, 18]. Moreover, it constitutes a crucial component of the innate defence system within the oral cavity [19, 20], possessing antimicrobial effector proteins, such as histatins and statherin [21]. Other contributors to host defence in the oropharynx include immunoglobulin A, cationic peptides, lysozyme, bactericidal/permeability-increasing protein, and mucins [19]. Although many salivary proteins have been characterised for their role in immune defence, it is unclear how the salivary proteome responds to acute oropharyngeal infection. Given the intrinsic variability of the salivary proteome [22], detecting subtle changes requires a well-controlled study design.

The CHIVAS-M75 trial [6], incorporated serial sampling of saliva before, during, and after infection with *emm75* GAS, providing a unique opportunity to examine the dynamic salivary proteome in response to infection. Here, we describe changes in immune-related and acute phase plasma proteins in saliva at key time points, illuminating features of the innate immune response at the site of infection.

## Methods

### Saliva collection

The CHIVAS-M75 trial is described in detail elsewhere [6]. Briefly, at the confirmed dose level, 20 healthy adults underwent controlled challenge using a swab to transfer an inoculum of *emm75* GAS onto the pharynx from a single-dose vial containing 10^5^ CFU in 1 mL of liquid animal-free media, with an estimated 10^4^ CFU adsorbed onto the swab and transferred. Saliva was collected during screening for GAS colonisation; the evening prior to challenge; 8-12 h post-challenge; and in 24-hour intervals following challenge until discharge 24 h after clinical diagnosis of pharyngitis (Figure 1A). Upon diagnosis, participants were commenced on intramuscular penicillin and oral rifampicin and discharged the next day. Most participants also provided follow-up saliva samples at 1-week, 1-month, and 3-months post-discharge. Unstimulated saliva was collected after a 30 min fast initiated with a tap water rinse. Participants rinsed their mouth again immediately prior to sampling and were instructed to produce 5-10 ml of saliva into a 50 ml Falcon tube on ice. With the exception of ‘evening prior’, all collections were conducted in the morning to minimise temporal variation upon glandular contributions [22]. Aliquots (1 ml) of unprocessed whole saliva were frozen immediately at −80 °C. The samples were not centrifuged to facilitate retention of mucin aggregates, though desquamated keratinocytes are also retained with this approach. Participants provided informed consent and were compensated for their involvement. The study was approved by the Alfred Hospital Human Research Ethics Committee (500/17).

**Fig 1.**
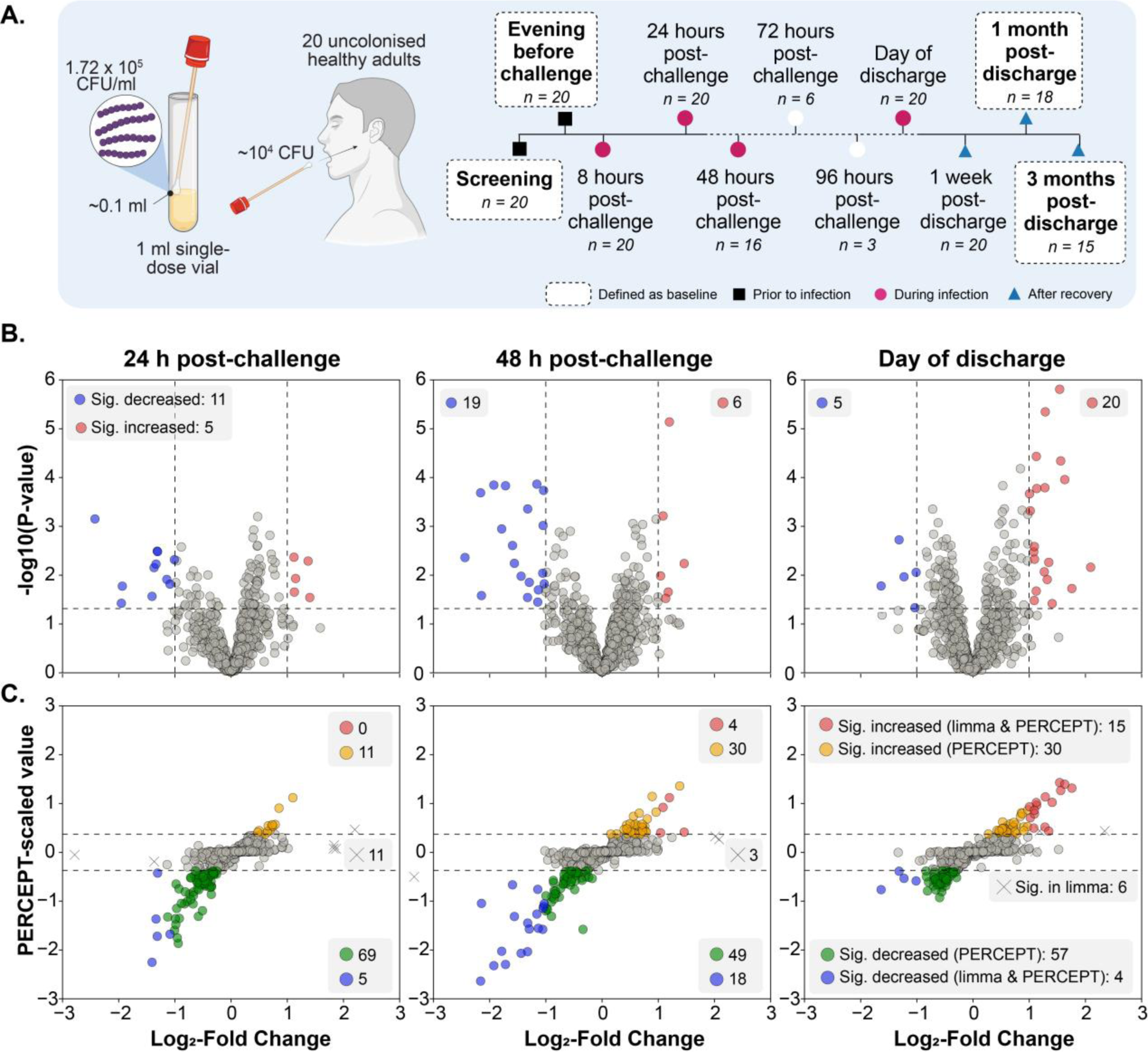
Significant proteins identified by PERCEPT during infection relative to baseline saliva. (A) Schematic representation of controlled streptococcal challenge and collection of saliva during CHIVAS-M75 trial. White-boxed time points indicated uninfected samples used as baseline. Differential abundance testing conducted via (B) limma and (C) PERCEPT with concordance represented by colour. Data are expressed as cohort mean of within-participant ratios comparing the average of baseline samples to 24 h post-challenge (n = 20); 48 h post-challenge (n = 16); day of discharge (variable timing; n = 20). For PERCEPT analysis, data-derived significance thresholds were calculated to exclude 95% of intrinsic variation (two-tailed; PERCEPT-scaled value ≤ −0.38 and ≥ 0.35; explained further in Supplementary Information (Figures S2, S5)).

### Sample processing

Saliva amount was normalised by volume and processed as adapted from previous works [23]. Protein from 100 µl of saliva was precipitated by overnight incubation in acetone. Pellets were dissolved in 8 M urea and reduced with 25 mM dithiothreitol followed by alkylation with 50 mM iodoacetamide. Proteins were digested overnight at 37 °C with 3 µg of sequencing-grade trypsin (Roche, Australia). Peptides underwent solid-phase extraction with C18 micro-columns prepared in-house. Detailed methodology including optimisation of sample processing approaches is provided in Supplementary Information.

### Peptide searches

Analysis of peptides by LC-MS was performed as described previously [24]. DIA mode was employed using an adapted BoxCar scheme [25]. Peptide data were searched against the reviewed human proteome (UniProtKB UP000005640) and annotated M75 challenge strain proteome (GenBank CP033621) using FragPipe v22.0 [26] in DIA mode with normalised label-free quantitation enabled. Further details are provided in the Supplementary Information.

To assess whether bacterial proteins originated from commensals, streptococcal proteins initially identified were screened against the Human Oral Microbiome Database (HOMD) [27] v11.02 using the “NCBI All Proteins” full-length protein search via Sequence Server 2.0.0 [28]. Additionally, precursor peptide sequences were searched in BLAST against a manually curated search space including genii represented from the HOMD search results (see Supplementary Methods).

### Data processing and statistical analyses

Common contaminants were removed after spectral data were searched [29, 30]. Keratin was considered a contaminant after confirming uniform distribution among samples (data not shown). To filter out rare or otherwise unreliably detected proteins, the following rule was applied: for inclusion in downstream analyses, proteins must be detected in at least three time points within at least three participants. Protein abundance at each time point was calculated relative to the average abundance across uninfected (‘baseline’) samples **(Figure 1A).** These ratios were log_2_-transformed where appropriate or further scaled using PERCEPT [31], a transformational framework enabling continuous, quantitative interpretation of p-values. This transformation follows conventional statistical comparisons; in this case, one-sample t-tests. Significant proteins were identified based on data-driven thresholding of the PERCEPT-scaled value at thresholds of 95% or 99% (Supplementary Information).

Protein abundance data were analysed by Ingenuity Pathway Analysis (QIAGEN IPA) version 122103623 (QIAGEN Inc.). Data were input as log_2_-fold-change and PERCEPT-scaled value with significance cut-offs for 95% variability as above. The core analysis was performed using default filters, limited to human data. Canonical pathway analysis was performed with Benjamini-Hochberg correction for multiple hypotheses.

Protein-protein interaction networks were constructed in Cytoscape 3.10.2 [32] using STRINGdb [33]. Streptococcal-human protein interactions were compiled from several host-pathogen interaction databases: HPIDB [34], APID [35], PHISTO [36], MENTHA [37] and manually appended from literature searches, including a recent detailed investigation of streptococcal interactions with human salivary and plasma proteins [38].

## Results

A total of 1315 proteins were detected across the cohort, including 71 matches to the M75 GAS proteome (Table S3). In all cases, these putative bacterial identifications occurred throughout the time-course, including in confirmed uninfected samples. BLAST-search of inferred full-length proteins against HOMD revealed high homology (Table S4) and subsequent search of detected peptides revealed complete sequence identity to non-*pyogenes Streptococci* and other commensals (Table S5). We therefore could not attribute these peptides to M75 GAS proteins and did not include them for further analysis, instead focusing on the human salivary proteome.

The salivary proteome exhibited highly variability both in the presence and absence of infection. Of 1244 human proteins detected in the cohort from at least one participant, 720 proteins were commonly detected in all 20 participants **(Figure S1)**. The abundance of individual proteins was also highly variable between participants and across sampling timepoints **(Figure S2)**. Indeed, no proteins exhibited stable abundance, defined as coefficient of variation <25%, in uninfected samples among all 20 individuals **(Table S2)**. Examination of technical and injection replicates demonstrated consistently low variation (Figure S3), affirming the source of variation to be purely biological. This intrinsic variability of the salivary proteome was reflected in a principal component analysis (PCA) **(Figure S4A)** where no clear clustering by time points was observed, though grouping by before, during, or after infection yielded separation of the 48 h post-challenge time point most extremely along both dimensions **(Figure S4B)**, followed by 24 h post-challenge, and day of discharge. These time points, which also correlated with peak clinical[6], cytokine, and cellular response [8], were chosen for further examination with respect to baseline, uninfected saliva (Figure 1A).

To examine differences in protein abundance, we used PERCEPT: an algorithm specifically developed for low-replicate, biologically noisy omics datasets [31]. PERCEPT in combination with data-driven thresholding (Figure S5) represents a less arbitrary, and more robust alternative to binary thresholding for identifying biological signals among highly variable data. PERCEPT identified a greater number of significantly different proteins than limma [39], though it did not uniformly lower detection rate as several proteins failed to retain significance when moving from limma to PERCEPT. The appropriateness of PERCEPT is further evident when compared to conventional testing with post-hoc correction (Figure S6; Supplementary Methods). Using PERCEPT, we identified 183 proteins that were significantly different to baseline in at least one of the three time points **(Figure 1B)**. The number and magnitude of significant differences was greatest at 48 h post-challenge. Four proteins were significant across all three time points **(Figure S7A)**, all decreased: Basic salivary proline-rich protein 1 (PRB1); Matrix Gla protein (MGP); Kinesin-1 heavy chain (KHC); ATP-dependent 6-phosphofructokinase platelet type (PKFP).

Hierarchical clustering was employed to examine whether changes in significant proteins were associated with clinical severity of pharyngitis (Table S6). Two participants (025 and 061) exhibited a broad increase of protein abundance at 48 h, a trend distinct from the rest of the cohort **(Figure 2)**. No grouping by pharyngitis grade was observed. Horizontal clustering, however, revealed several subsets of consistently enriched proteins at 48 h, including a group of primarily plasma proteins, including inflammation-related and acute phase proteins such as complement component 3 (C3), c4b-binding protein (C4BP), fibrinogen (FGA, FGB, FGG), ceruloplasmin (CP), alpha-1-antitrypsin (A1AT). Bootstrap resampling yielded significance (AU > 0.95) of this cluster. Conversely, clusters of salivary proteins were suppressed across the cohort, including the family of basic proline-rich salivary proteins and other oral homeostasis-associated proteins. From these observations, it was postulated that plasma- and saliva-origin proteins at 48 h post-challenge exhibited different abundance trajectories with respect to baseline; namely that plasma-origin proteins increased while salivary proteins decreased. This hypothesis was accepted both when considering all proteins and also when restricted to significant proteins (Mann-Whitney U test, p < 0.0005, Figure S7).

**Fig 2.**
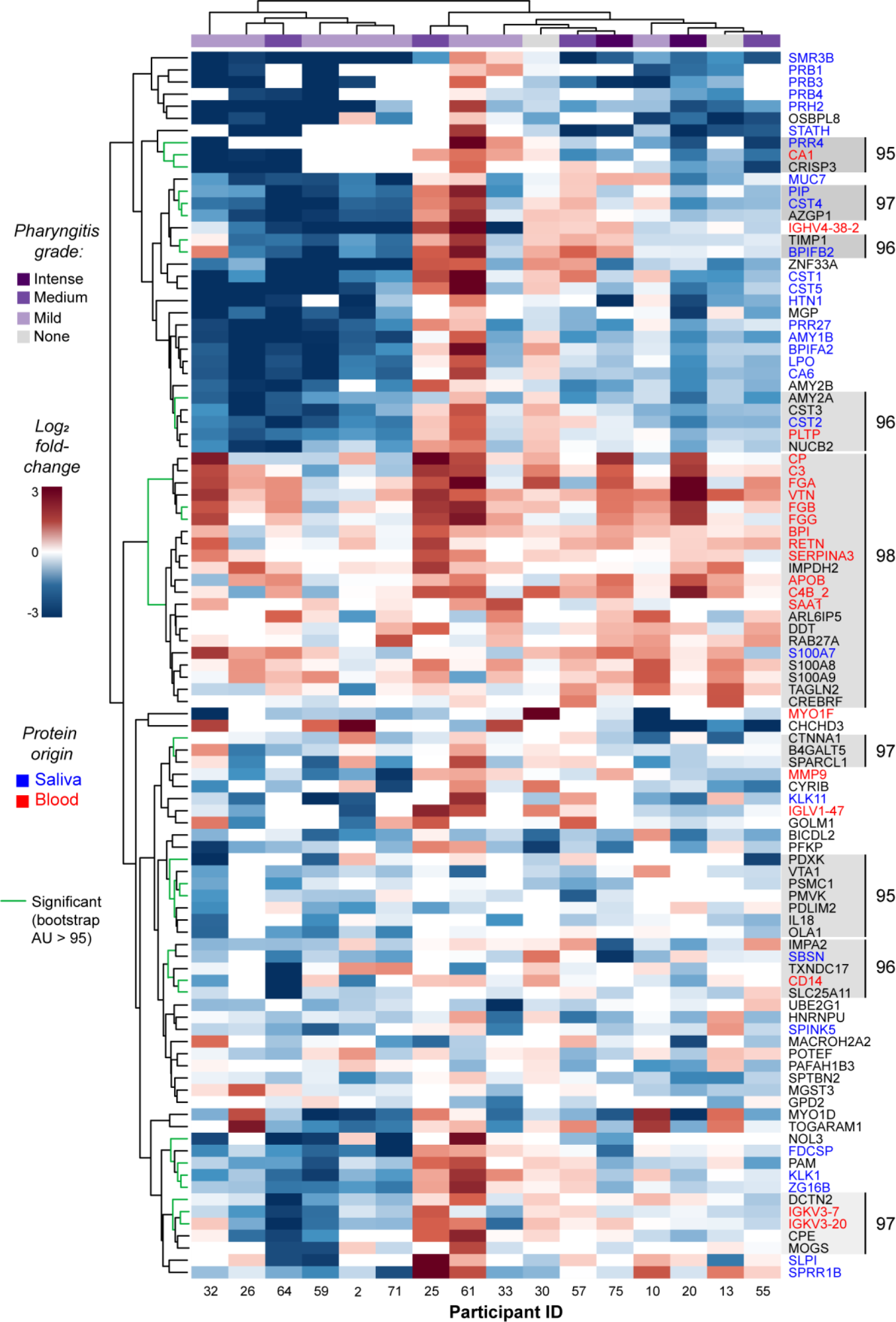
Protein abundance at 48 h post-infection relative to baseline. Protein quantity is expressed relative to baseline per participant (n = 16). Grade of pharyngitis is displayed as reported elsewhere [6]. Protein labels are coloured by predicted tissue origin determined by literature search and comparison to the Human Salivary Proteome Database [40] and Human Protein Atlas [41]. Both axes are hierarchically clustered using Ward’s method and Euclidean distancing with zero-filling of missing ratios. Branch reliability of protein dendogram was assessed by bootstrap resampling with significance indicated with green highlights at approximately unbiased (AU) values >95%. Grey shading indicates the significantly grouped proteins at highest hierarchy with AU values reported adjacently.

After observing broad changes at the level of protein origin, we then examined infection-related changes among individual proteins. To identify the most extreme and consistent changes, we employed a protein detection filter of 50% (n=10) within the cohort at a given time point and a more conservative threshold to exclude 99% of intrinsic sample variability. This approach yielded 33 consistently altered proteins. The abundance patterns and dynamics of these proteins were associated with their canonical origin in plasma or saliva. All 12 plasma-derived proteins increased in abundance post-infection and were associated with immune functions, particularly the acute phase response **(Table 1)**. In contrast, most proteins that decreased were of salivary origin and many lacked known immune functions **(Table S7)**. Intriguingly however, several salivary proteins with known antimicrobial activity also decreased in abundance including BPI fold-containing family A member 2, lactoperoxidase, cystatin D, histatin 1, and mucin-7. Plasma and salivary proteins differed temporally as well. Salivary antimicrobial proteins decreased at 24 and 48 h post-infection, but not at discharge, whereas acute phase and complement system-related proteins were exclusively significant at discharge. At this threshold, no proteins were significant across all three post-infection time points **(Figure S4B)**.

**Table 1.** List of consistently altered immune-related proteins at during infection. Proteins are included if surpassing the 99% data-derived threshold in at least one time point and detected in at least 10 participants. + indicates increase relative to baseline, - indicates decrease. A full table without filtering is provided in Supplementary Information. Immune and non-immune functions are described generally as per literature review. Some categories overlap with proteins having several functions.

| Immune-related function | Gene | +/- | Protein | Non-immune function | Typical of: | Time point(s) |
| --- | --- | --- | --- | --- | --- | --- |
| Acute phase protein | ORM1 | + | Alpha-1-acid glycoprotein 1 | Transport | Plasma | Discharge |
|  | CP | + | Ceruloplasmin | Copper transport | Plasma | Discharge |
|  | HP | + | Haptoglobin | Haemoglobin binding | Plasma | Discharge |
|  | HPX | + | Hemopexin | Heme binding | Plasma | Discharge |
| Antimicrobial | BPIFA2 | - | BPI fold-containing family A member 2 | - | Saliva | 24 h, 48 h |
|  | CST5 | - | Cystatin-D | Cysteine protease inhibitor | Saliva | 24 h, 48 h |
|  | CST2 | - | Cystatin-SA | Thiol protease inhibitor | Saliva | 48 h |
|  | HTN1 | - | Histatin-1 | Wound healing | Saliva | 24 h, 48 h |
|  | LPO | - | Lactoperoxidase | Thiocyanate oxidiser | Saliva | 24 h, 48 h |
|  | MUC7 | - | Mucin-7 | General oral homeostasis | Saliva | 48 h |
| Cell death | GSDMA | + | Gasdermin-A | Pyroptosis | Neither | Discharge |
| Immunity | C3 | + | Complement C3 | - | Plasma | Discharge |
|  | CFB | + | Complement factor B | - | Plasma | Discharge |
| Immuno-modulatory | SERPINA3 | + | Alpha-1-antichymotrypsin | Protease inhibitor | Plasma | Discharge |
|  | SERPINA1 | + | Alpha-1-antitrypsin | Protease inhibitor | Plasma | Discharge |
|  | VTN | + | Vitronectin | Adhesion molecule | Plasma | 48 h |
|  | CST3 | - | Cystatin-C | - | Both | 48 h |
|  | PLTP | - | Phospholipid transfer protein | Cholesterol transport | Plasma | 48 h |
|  | SAA1 | - | Serum amyloid A-1 protein | Lipid metabolism | Plasma | Discharge |
| Inflammation | FGA | + | Fibrinogen alpha chain | Clotting | Plasma | 48 h, Discharge |
|  | FGB | + | Fibrinogen beta chain | Clotting | Plasma | 48 h, Discharge |
|  | FGG | + | Fibrinogen gamma chain | Clotting | Plasma | 48 h, Discharge |

After identifying cohort-wide changes in protein abundance, we next examined these proteins levels within individual participants. Acute phase, innate immune, and immunomodulatory proteins increased in most participants **(Figure 3)**. Acute phase, innate immune, and immunomodulatory proteins exhibited consistent increases across the cohort **(Figure 3)**. This trend was noted both in high-abundance proteins detected commonly, such as complement C3, and in proteins rarely detected in uninfected saliva, such as alpha-1-antichymotrypsin. An association between these infection-related proteins and degree of clinical pharyngitis was hypothesised, but not supported by hierarchical clustering as increased levels were apparent even in non-pharyngitis participants.

**Fig 3.**
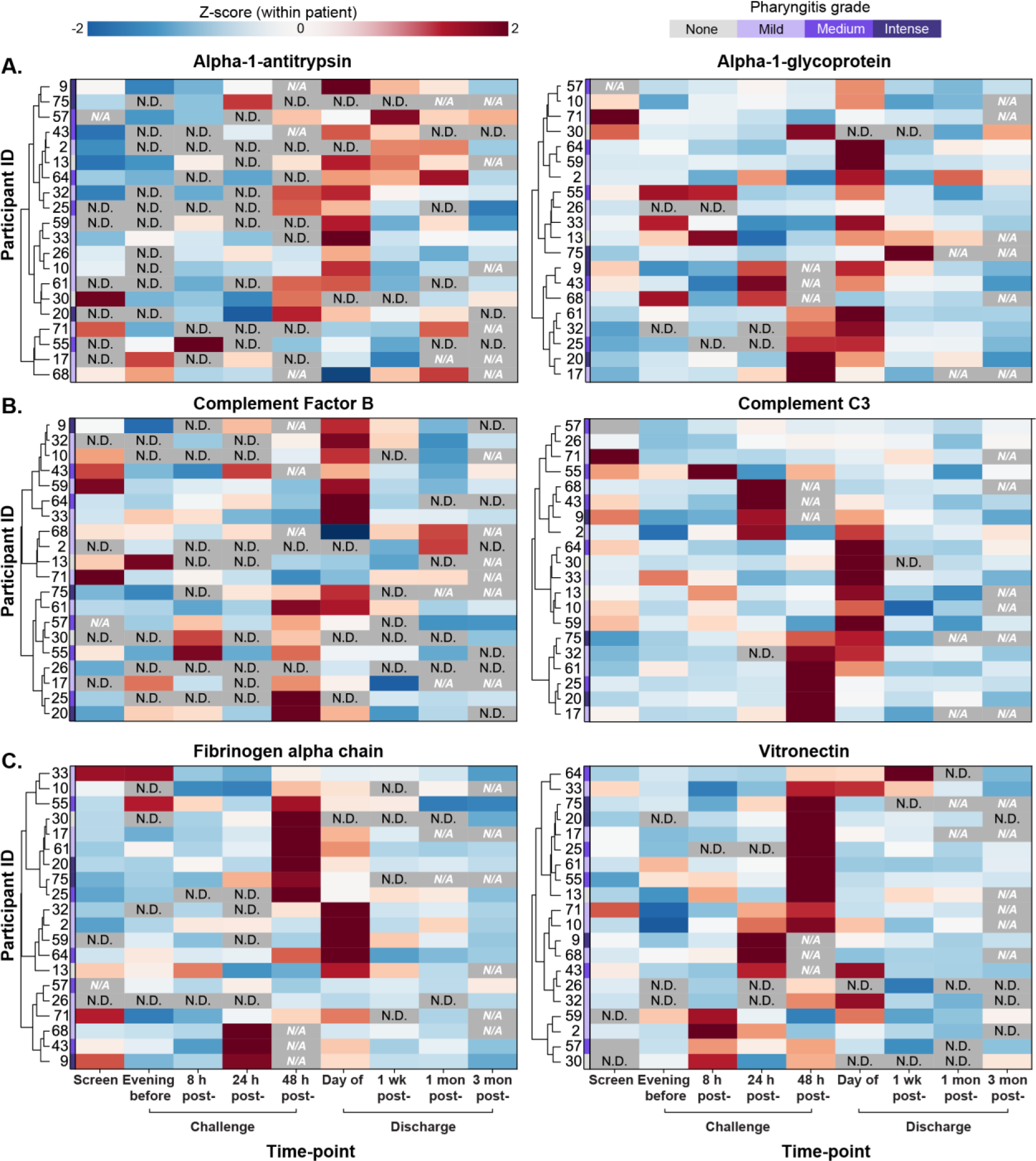
Dynamics of selected acute phase proteins (A), innate immune proteins (B), and immunomodulatory proteins (C). Z-scores were calculated within-participant per protein. Participants are hierarchically clustered using Ward’s method and Euclidean distancing. Missing data were imputed for non-detection events using K-Nearest Neighbours method for clustering purposes only but are displayed as N.D. for undetected proteins at a given time point, and N/A for participants who were not sampled at that time point. Grade of pharyngitis is displayed as reported elsewhere [6].

Participants 013 and 030 were the only two individuals not diagnosed with clinical pharyngitis, presenting a unique opportunity to qualitatively examine whether observed proteomic changes were driven by bacterial colonisation or symptomatic pharyngitis. The former was heavily colonised without symptoms, signs or systemic inflammation whereas the latter was uncolonised. Strikingly, trajectories of significant proteins in the subclinical-colonised participant (013) closely tracked the pharyngitis cohort, whereas the the uncolonised participant (030) exhibited notable deviation **(Figure 4)**.). This discrepancy between the two non-pharyngitis participants was observed in general salivary proteins (Figure 4A), immune-related salivary proteins (Figure 4B), and acute phase proteins (Figure 4C).

**Fig 4.**
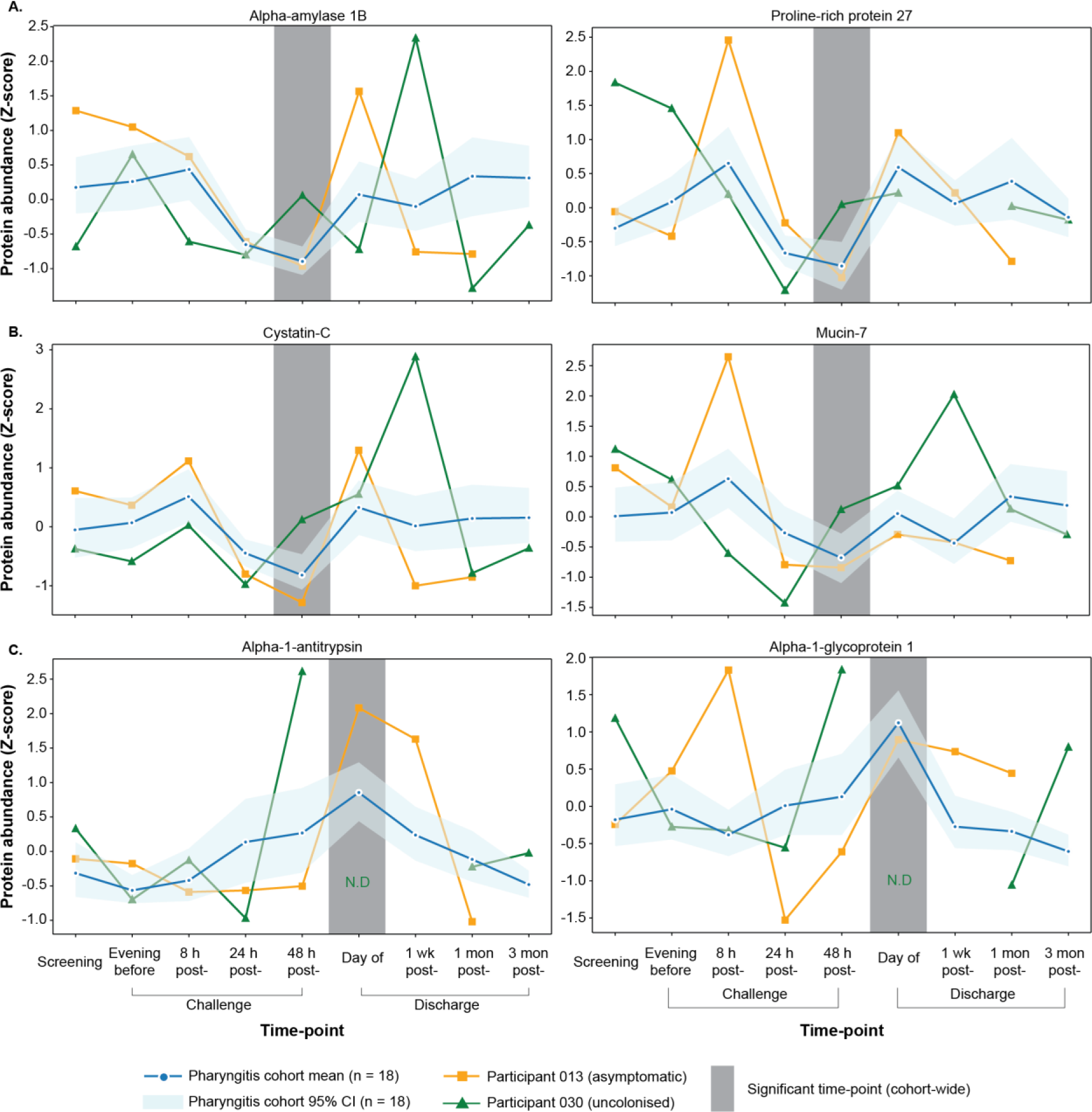
Lineplots of selected general salivary proteins (A), immune-related salivary proteins (B), and acute phase plasma proteins (C). Pharyngitis-positive cohort is represented as mean with 95% confidence interval (n=18). Participants 013 and 030 are plotted separately. Protein abundance is represented as within-participant-calculated Z-scores. The blue asterisk denotes the time point(s) in the pharyngitis-positive cohort at which abundance is significantly different to baseline as described earlier; N.D. denotes non-detection of the given protein at the significant time point.

After observing changes at the level of individual proteins, pathway-level changes were examined at each time point (Figure 5A-C). Relative to baseline, significant proteins were enriched for LXR/RXR activation and acute phase response signalling pathways at day of discharge. These two pathways were also elevated at 48 h, in addition to DHCR24 signalling and the coagulation system **(Figure 5D)**. No enrichment of any pathway was detected at 24 h. The protein-protein interaction network for these proteins supported this finding, revealing a striking increase in the abundance of a cluster of plasma proteins, including members of the acute phase response **(Figure 6)**. In stark contrast, salivary proteins clustered separately and were decreased relative to baseline. All known streptococcal interactors of the altered proteins were also included in the network **(Figure 6; purple boxes)**, predominantly interacting with the cluster of elevated plasma proteins. This visualisation is consistent with host ligands of streptococcal proteins increasing in saliva at 48 h.

**Fig 5.**
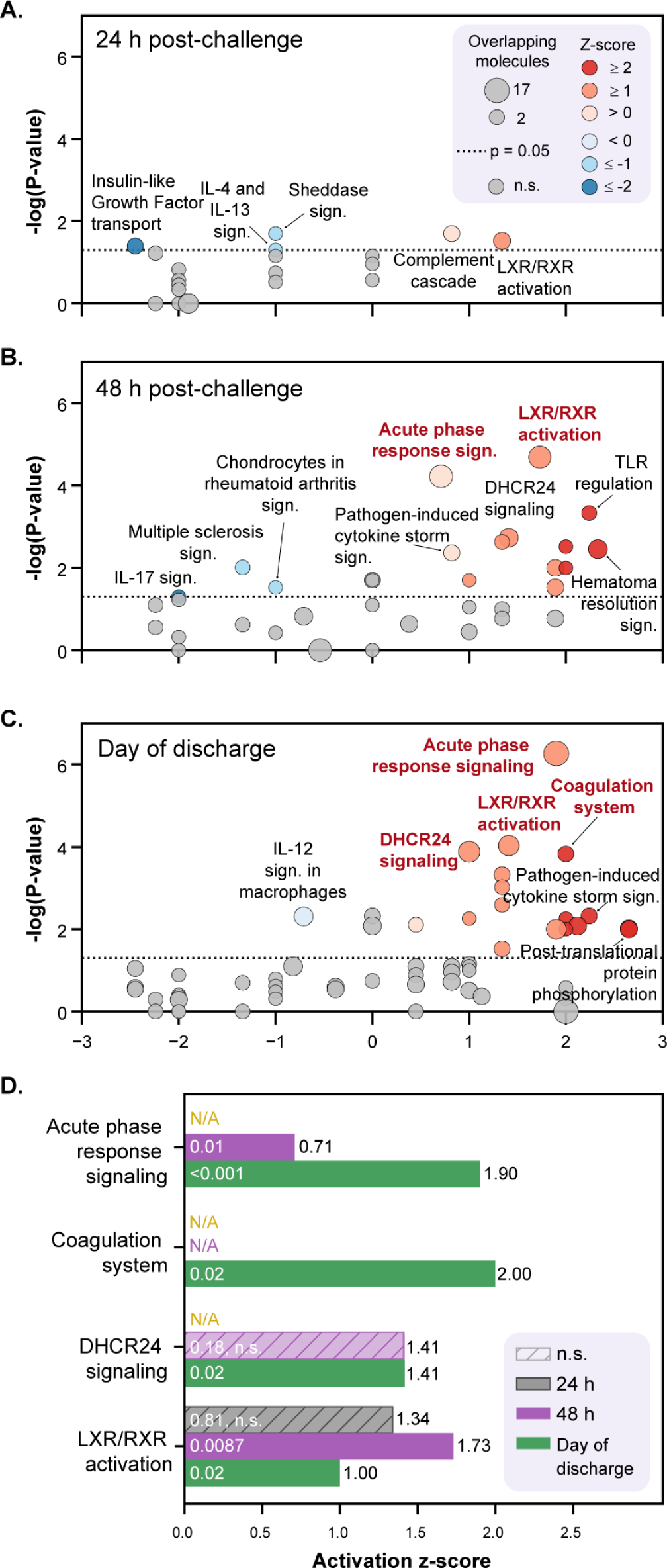
Pathway-level enrichment across the post-challenge time-course. Canonical pathways identified as significant (p<0.05) in Ingenuity Pathway Analysis at 24 h post-challenge (A), 48 h (B), and day of discharge (C). Four pathways remain significant after Benjamini-Hochberg (BH) correction represented in bold and summarised in (D), with adjusted p-values reported in white-text within the bar and activation z-score reported adjacently. “N/A” represents pathways that were not identified at that time point. Proteins identified as significant using a 95% PERCEPT-scaled value threshold were evaluated against all proteins detected experiment-wide, with log_2_-fold change values considered as the quantitative metric. Pathway data are available in Table S8.

**Fig 6.**
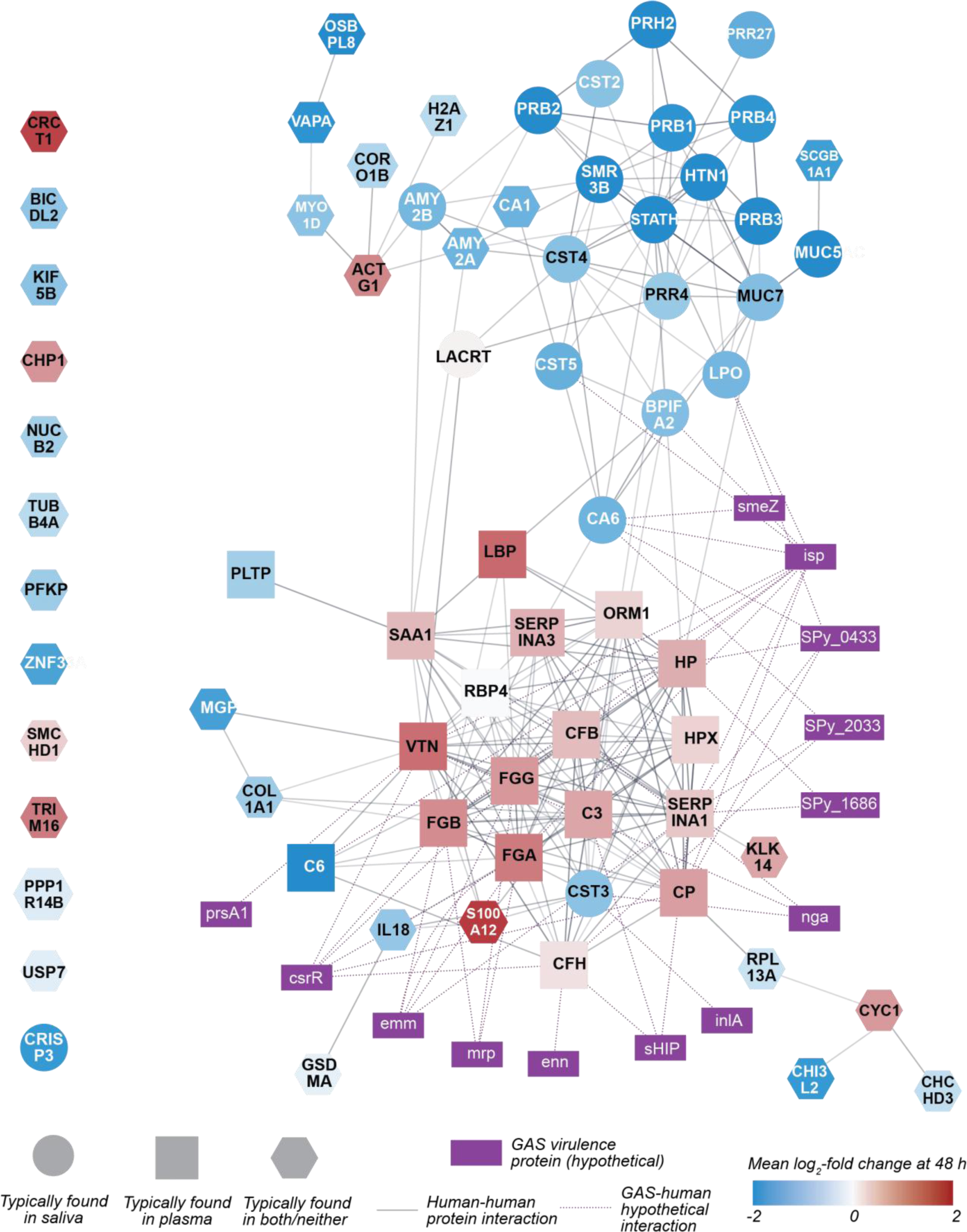
**Protein-protein interaction network of significant proteins and known (hypothetical) GAS virulence factors,** colour-mapped by relative abundance at 48 h. Purple rectangular nodes represent known streptococcal interactors and purple dotted lines represent evidenced streptococcal-human protein-protein interactions. Grey lines between human nodes represent evidenced or predicted protein-protein interactions with darker lines representing higher StringDB interaction score. Unclustered human proteins are listed on the left. Node positioning has been slightly modified for aesthetic purposes.

## Discussion

This study provides the first comprehensive characterisation of the dynamic protein response to GAS pharyngitis, revealing both temporal and compartmental shifts at the site of infection during acute illness. Our findings indicate that GAS infection triggers a consistent change of the salivary proteome, driving a decrease of antimicrobial salivary proteins and concurrent influx of immune-related and acute phase plasma proteins, many of which are targets of known GAS virulence factors. Moreover, the differential protein responses observed between participants with clear acute pharyngitis and asymptomatic colonised and uncolonised individuals hints at potential mechanisms of host resistance to GAS infection. This work provides the first direct evidence of a consistent, infection-driven shift in the salivary proteome, deepening our understanding of the mucosal response to GAS infection.

Saliva is heralded as an attractive sample for diagnostic and prognostic molecular markers due to the ease of collection and low dynamic range compared to serum [42–44]. It is also of particular relevance for acute illness in the oropharynx. Saliva from the CHIVAS-M75 trial has previously undergone targeted analysis revealing differential IgA responses between pharyngitis positive and negative groups [10] and elevated cytokine profiles [8]. Mass spectrometry facilitates a broad, untargeted, discovery-focused approach to complement such analyses. Using BoxCar DIA, which leverages advanced mass spectrometric hardware and neural-network informed search strategies [25, 45], we identified a total of 1244 distinct proteins across the cohort, exhibiting superior coverage than previous DDA analyses of saliva identifying 100-400 [46–50] or 400-600 [51–53] proteins. To our knowledge, this is the first study to analyse the salivary proteome using boxcar DIA mass spectrometry, allowing unprecedented, unbiased biological insights with minimal sample handling.

Pharyngitis drove a change in the salivary proteome, with the greatest changes occurring at 48 h post-challenge. This timing corresponds to peak pharyngitis in most participants as measured by bacterial load, markers of systemic inflammation in serum, and other clinical parameters [6]. Of the significant proteins detected consistently in the cohort post-infection, the elevated fraction entirely comprised plasma proteins. Plasma proteins may enter saliva via gingival crevicular fluid, which increases substantially during oropharyngeal inflammation [54, 55] or by passive diffusion across epithelial and endothelial barriers of the oral mucosa [43] Notably, albumin, the primary component of the plasma proteome, is not elevated during peak infection, suggesting some degree of specificity in leakage. Indeed, all the proteins which were elevated have immune-related functions, including the acute phase response. This selective increase may reflect systemic upregulation of these acute phase proteins or local host-pathogen interactions, or a combination of both, though these mechanisms cannot be resolved by proteomics data alone.

Acute phase reactants respond non-specifically to trigger systemic production of inflammatory products and include cytokines and C-reactive protein (CRP), complement-related proteins, haptoglobin, ceruloplasmin, serum amyloid A, alpha-1-acid glycoprotein (AGP), and fibrinogen [56]. Neither cytokines nor CRP were among the consistently increased acute phase proteins detected in saliva (Table 1). Although serum CRP was universally elevated in the cohort during infection [6], saliva and serum CRP do not consistently correlate [57]. Previous work found increased cytokines at 24 h and 72 h [8], which was not apparent in our untargeted analysis. This likely reflects the low abundance of these analytes in saliva, for which targeted analyses retain a sensitivity advantage over untargeted proteomics. Other acute phase reactants were consistently enriched during infection and this effect was detectable irrespective of typical abundance in saliva. For example, fibrinogen was consistently detected in participants across time points and can be considered of at least moderate abundance in saliva. A1AT, however, was undetectable in half of the cohort on the evening prior to infection but detected in 17 of 20 participants on the day of discharge at elevated levels. Acute phase reactants were detected even for usually low-abundance proteins.

We observed distinct temporal dynamics between proteins and between participants. For instance, fibrinogen exhibited an earlier and more transient increase compared to the more persistent elevation of C3 and AGP. In most participants, acute phase reactants peaked on the day of discharge, when typical salivary proteins had returned to baseline levels. While participants with and without pharyngitis showed similar protein response patterns, the uncolonised participant (030) exhibited markedly different responses, particularly in acute phase proteins. This distinction suggests that bacterial colonisation alone, independent of symptomatic pharyngitis, may be sufficient to drive proteomic changes in saliva. Interestingly in the colonised asymptomatic participant (013), there was an increase in the antimicrobial proteins MUC7 and A1GP at 8 h post-challenge not seen in other participants. Though limited by sample size, these differences suggest potential mechanisms of host resistance to GAS infection. The subtlety and complexity of this putative response is reflected in previous studies of saliva from this model [8, 10].

GAS is a highly adapted human-restricted pathogen that has co-evolved to exploit the human inflammatory response in its pathogenesis [58]. Among proteins significantly increased in this study, many were plasma proteins known to be targets of streptococcal virulence factors. Although *in vitro* and biochemical research has previously proposed the exploitation of target human proteins by GAS, our study is the first to demonstrate the enrichment of these at the site of infection in human pharyngitis, suggesting further their relevance *in vivo*. Fibrinogen plays a crucial role in host defence against oral pathogens, as salivary factors enhance its capacity to entrap bacteria in fibrin clots [59]. However, GAS is known to have evolved beyond this defence by degrading fibrinogen and other clot-related components [60]. Moreover, fibrinogen-binding by GAS is well-established to confer protection against host immunity, including complement deposition [61, 62]. Notably, all three chains of fibrinogen and elements of the complement system such as complement factor B, C3, and C4BP were all increased during infection. Similarly, vitronectin, which is bound [63] and degraded [60] by GAS, was increased in all participants for at least one time point. Vitronectin recruitment by pathogens has been implicated in complement evasion [64], although this has not previously been explored for GAS.

The increase in plasma proteins was accompanied by a marked decrease in salivary proteins. Notably, every significant protein identified as salivary was decreased rather than enriched. While many of these lack known immune function, there were several examples of resident antimicrobial proteins such as cystatin-D and mucin-7 decreasing during infection. These saliva proteins are considered first-line defences [19], more likely acting to deter colonisation rather than combat active infection. Mucosal transcriptomic analyses, outside the scope of this study, could conclusively determine whether decreased protein abundance is a consequence of decreased transcription, ‘dilution’ from infiltrating plasma proteins, or consumption in the overall local antimicrobial process. Nonetheless, in the context of a highly variable environment, we have identified a consistent, marked, and biologically relevant change in the salivary proteome in response to GAS infection.

The current study has a number of limitations. Principally, this is a small cohort with limited statistical power, and findings may not be generalisable to all populations or indeed all strains of GAS. Salivary flow rate was not measured, meaning relative abundance changes cannot be fully decoupled from dilution effects. Finally, the pathway and network analyses conducted here are exploratory in nature and should be treated as hypothesis-generating rather than mechanistically conclusive. These limitations are offset by the unique advantages intrinsic to controlled human infection models, which eliminates common confounders in observational studies, such as variable infectious dose and strain, unknown time of infection, and pre-existing infection or carriage. The depth of proteomic coverage and subsequent application of data-driven thresholding ensured that the identified changes reflect robust, biologically meaningful signals despite the inherent variability of saliva.

The experimental human pharyngitis model provides a remarkable opportunity for the interrogation of host-pathogen biology. In our detailed analysis of the dynamic salivary proteome during GAS pharyngitis, we demonstrated a consistent, infection-driven depletion of resident antimicrobial salivary proteins alongside selective enrichment of acute phase plasma proteins in the oropharynx, many of which are established targets of streptococcal virulence factors. These insights advance our understanding of the dynamic mucosal environment in GAS pharyngitis and affirm the value of salivary proteomics as an important endeavour in understanding pathogenesis in the human oropharynx.

## Data and code availability

Mass spectrometry data has been deposited to the ProteomeXchange Cosortium via PRIDE [65] with the dataset identifier PXD068717. Custom Python analysis scripts used in this study are available from Zenodo via https://doi.org/10.5281/zenodo.14917970.

## CRediT author statement

*Conceptualisation*: N.P., A.E.D., M.L.S-S.; *Methodology:* A.D.I., C-H.L., A.E.D., N.P., M.L.S-S.; *Formal analysis:* A.D.I., D.C.; *Investigation:* A.D.I., P.K.D., A.E.D.; *Resources:* J.O., K.A., A.S., D.C., A.E.D., M.L.S-S.; *Data curation:* A.D.I., D.C.; *Writing – Original Draft:* A.D.I, *Writing – Review & Editing:* A.D.I., D.S., M.L.S-S., A.E.D.; *Visualisation:* A.D.I., D.C.; *Supervision:* D.S., A.E.D., M.L.S-S.; *Project administration*: J.O., K.A., A.S., A.E.D, M.L.S-S.; *Funding acquisition:* A.E.D., N.P., M.L.S-S.

## Acknowledgements

The authors thank Hannah Frost for critical reading, Queensland Cyber Infrastructure Foundation for bioinformatics support, and all participants and staff involved with the CHIVAS trial. This study was funded by the National Health and Medical Research Council, Australia. A.D.I was supported by the Australian Government Research Training Program. No authors declare any conflict of interest.

## Supplementary Information

### Methods

#### Optimisation of methodology

Several elements of sample preparation and search strategies were initially assessed using aliquots of saliva from donors outside of the CHIVAS trial and evaluated primarily by two metrics: co-efficient of variation (CV) between technical replicates and number of protein group identifications (IDs). Acetone-precipitated protein yielded 5.1% more identifications than treatment of otherwise unprocessed dried saliva. Though this difference did not reach statistical significance (p = 0.12, paired t-test), CV improved from 33% to 19%. Inclusion of solid-phase extraction (SPE) with in-house C18 micro-columns (described below) improved chromatographic robustness and reproducibility (data not shown) without affecting IDs (p = 0.78). IDs among C18-enriched peptides derived from acetone-precipitated material did not improve when using a more lenient DDA search strategy (Proteome Discoverer 2.5; up to 4 missed cleavages, up to 4 variable modifications) relative to a stricter search space (up to 2 missed cleavages, up to 2 variable modifications). Analysis of corresponding spectra generated from DIA mode yielded greater proteomic depth in Spectronaut (87% more IDs) and DIA-NN 1.8 (145% more IDs).

Following data acquisition from CHIVAS peptides, Fragpipe v23 with built-in DIA-NN 2.0 was employed after comparisons to standalone DIA-NN 1.8 generated more reproducible peptide identification between biological replicates (data not shown).

#### Assessment of technical variation

Reproducibility of sample preparation and LC-MS analysis were examined by technical and injection replicates, respectively. A single CHIVAS saliva sample with excess volume yielded five aliquots which independently underwent the complete laboratory workflow. One of these aliquots was analysed repeatedly in five independent injections. These comparisons were also performed upon a pooled saliva sample comprised of eight randomly selected individuals. Injection order was randomised. Protein CVs were calculated on DIANN-searched DIA data as per [66] and replicate concordance was interrogated by Pearson correlation.

#### Sample preparation

Saliva samples were thawed on ice for 2 h. Protein from 100 µl of homogenised was precipitated overnight by incubation at −20 °C with 900 µl of acetone. Protein pellets obtained by centrifugation at 14,000 *× g* for 30 min at 4 °C were resuspended by bath-sonication in 10 µl of 8 M urea for 5 min, followed by addition of 10 µl water, and further sonication for 10 min. Disulphide bonds were reduced by incubation with 25 mM dithiothreitol for 3 hr at 60 °C, followed by alkylation with 50 mM iodoacetamide incubated at room temperature for 30 min, protected from light. Denatured, reduced, and alkylated protein sample was digested overnight at 37 °C with 3 µg of sequencing-grade trypsin (Roche, Australia; #11418475001) in a total reaction volume of 50 µl of 50 mM ammonium bicarbonate, pH 8. Trypsin reaction was stopped by heat-inactivated at 70 °C for 10 min and samples were dried in a Savant^TM^ SpeedVac vacuum concentrator (Thermo Fisher Scientific) in preparation for enrichment by SPE. C18 micro-columns were prepared by excision of Empore^TM^ C18 (Merck, Australia; #66883-U) material into 200 µl pipette tips C18 columns were equilibrated with two washes of 80% (v/v) acetonitrile (ACN) in 0.1% (v/v) aqueous trifluoroacetic acid (TFA), followed by two washes with 0.1% TFA. All spins were performed at maximum speed for 30-60 s on a benchtop minicentrifuge at room temperature. Dried peptides were resuspended in 0.1% TFA and loaded into micro-columns. After centrifuging, flowthrough was reloaded and washed with two column volumes of 0.1% TFA. Peptides were eluted with two column volumes of 80% ACN, 0.1% TFA and dried in a vacuum concentrator.

#### Liquid chromatography and mass spectrometry

Peptide separation and mass spectrometry analyses were performed using an UltiMate^TM^ 3000 RSLCnano HPLC system coupled to an Orbitrap Fusion^TM^ mass spectrometer via a Nanospray Flex^TM^ ion source (all components from Thermo Fisher Scientific, USA). Peptides were initially loaded onto an Acclaim^TM^ PepMap^TM^ 100 C18 pre-column (75 μm x 20 m; Thermo Fisher Scientific) and separated on a MonoCap C18 Nano-flow column (0.1 × 150 mm; GL Sciences, USA) maintained at 45 °C. Separation was achieved using a 130-minute gradient of buffer A (0.1% formic acid) and buffer B (80% acetonitrile with 0.1% formic acid) at a flow rate of 300 nL/min.

Samples were analysed using a BoxCar data-independent acquisition scheme. MS1 scans were acquired from m/z 350-1650 at 120,000 resolution, with 24 variable-width segments distributed across three multiplexed targeted SIM scans isolating eight segments each. The normalized AGC target was set to 200% per segment with a 246 ms maximum injection time. The MS2 segments maintained the same segmentation strategy with a 1-m/z overlap between adjacent windows. MS2 spectra were recorded at 30,000 resolution with 2000% AGC target and 60 ms maximum injection time. Fragmentation was performed using stepped HCD at collision energies of 22%, 27%, and 32%.

#### DIA analysis and post-processing

Data files were searched in Fragpipe v 22.0 [26] using a modified version of the “DIA_SpecLib_Quant” workflow with spectral library generation, using the reviewed human proteome (Uniprot KB UP000005640) with decoys. Peptide length was set to 7-50 residues, allowing up to one missed tryptic cleavage. Precursor and fragment mass tolerances were set to 10 and 20 ppm, respectively. Cysteine carbamidomethylation was enabled as a fixed modification whereas methionine oxidation, and N-terminal methionine removal, and N-terminal acetylation were allowed as variable modifications. Quantitation was handled by DIA-NN 2.0 within Fragpipe environment with default settings including match-between-runs (MBR) and without imputation. Quantitation data were exported from DIA-NN and manipulated in Python. As per default settings, results are filtered out at 1% FDR at both protein and peptide level. Prior to analysis in Fragpipe, .raw files were converted to mzml in MS Convert 3.0 [67]. Technical (injection) replicates were not included here following demonstration of technical reproducibility described above.

CHIVAS saliva was examined for GAS proteins in a separate analysis against a combined search space of the reviewed human (Uniprot KB UP000005640) and M75 translated proteome (GenBank CP033621), with MBR disabled to prevent false positive carry-over detections. To assess whether these proteins arise from M75 GAS or oral commensals, bacterial proteins identified within the M75 proteome were screened against HOMD at a full-length protein level. Subsequently, peptide sequences attributed to M75 proteins in the original DIA search were also screened in BLAST against a curated search space generally including genii of oral commensals and specifically including genii represented in the protein-level search.

#### Data-driven thresholding of scaled fold-change values

Thresholds to isolate biologically relevant proteins were determined by a data-driven approach [31]. Following one-sample t-tests, PERCEPT-scaled values were calculated between randomly selected pairs of baseline time points within individual participants (e.g. baseline vs 3-months post-discharge). The mean of these ratios across all participants was then used to generate a distribution of log_2_-fold change representing estimate the natural biological variation in the absence of any experimental intervention (i.e. challenge). From this distribution, thresholds capturing 95% of the variance (i.e. at the 2.5^th^ and 97.5^th^ percentile) were determined. To reduce the impact of random time point selection (Figure S3), this process was repeated 100 times. The mean lower and upper values of these 100 iterations were used as the final thresholds. These thresholds therefore bound the expected natural variation in the system, beyond which changes in protein abundance are considered significant.

## Supplementary tables

**Table S1.** Summary of CHIVAS- and CHIPS*-related publications.

| First author, year | Primary focus |
| --- | --- |
| Osowicki, 2021 [6] | Dose-finding, clinical validation |
| Fabri, 2021 [7] | Molecular quantification of colonisation |
| Anderson, 2022 [8] | Immune profiling, transcriptomics |
| Indraratna, 2024 [9] | Diagnostic saliva qPCR |
| Osowicki, 2024 | Mucosal and systemic antibody responses |
| Enkel, 2024 [11] | Transmission dynamics (CHIPS)* |
| Hla, 2025 [12] | Penicillin pharmacokinetics (CHIPS)* |
| Fryer, Pitt, 2026 [13] | Glycan-specific B cell responses |
| Lepletier, 2025 (preprint) [14] | Longevity of IgG responses |

**Table S6.** Grouping of pharyngitis severity based on visual evaluation by trained physicians [6]. Participants who did not meet clinical diagnosis are categorised as ‘None’. Participants are organised from earliest to latest diagnosis. *Denotes participants who were discharged before 48 hours and therefore not represented in 48 h time point.

| *Pharyngitis severity* | *Participant ID(s)* |
| --- | --- |
| None | 013, 030 |
| Mild | 068*, 002, 017, 033, 059, 061, 071, 032, 010, 026 |
| Medium | 043*, 025, 064, 055, 057, |
| Intense | 009*, 020, 075 |

**Table S7.** List of significant non-immune proteins at 99% PERCEPT threshold. (v ≤ −0.73 and v ≥ 0.68). Protein functions are described generally as per literature review. All listed proteins were decreased at the listed time point(s).

| **Gene** | **Protein** | **Function** | **Time point(s)** | **Origin** |
| --- | --- | --- | --- | --- |
| AMY1B | Alpha-amylase 1B | Carbohydrate digestion | 24 h, 48 h | Saliva |
| AMY2A | Pancreatic alpha-amylase | Carbohydrate digestion | 24 h, 48 h | Saliva |
| AMY2B | Alpha-amylase 2B | Carbohydrate digestion | 24 h, 48 h | Saliva |
| BICDL2 | BICD family-like cargo adapter 2 | Intracellular transport | 48 h | Neither |
| CA1 | Carbonic anhydrase 1 | General homeostasis | 48 h | Neither |
| CA6 | Carbonic anhydrase 6 | General oral/dental homeostasis | 24 h, 48 h | Saliva |
| CRISP3 | Cysteine-rich secretory protein 3 | Reproductive immunity | 24 h, 48 h | Saliva |
| CST4 | Cystatin-S | General oral/dental homeostasis | 48 h | Saliva |
| MGP | Matrix Gla protein | Bone metabolism | 24 h, 48 h | Neither |
| NUCB2 | Nucleobindin-2 | Calcium homeostasis | 24 h | Both |
| OSBPL8 | Oxysterol-binding protein-related protein 8 | Lipid transport | 24 h, 48 h | Neither |
| PFKP | ATP-dependent 6-phosphofructokinase, platelet type | Glycolysis | 48 h | Neither |
| PRB1 | Basic salivary proline-rich protein 1 | General oral/dental homeostasis | 24 h, 48 h | Saliva |
| PRB3 | Basic salivary proline-rich protein 3 | General oral/dental homeostasis | 24 h, 48 h | Saliva |
| PRB4 | Basic salivary proline-rich protein 4 | General oral/dental homeostasis | 24 h, 48 h | Saliva |
| PRR27 | Proline-rich protein 27 | Unknown | 24 h, 48 h | Saliva |
| RPL13A | Large ribosomal subunit protein uL13 | Ribosomal | Discharge | Neither |
| SMR3B | Submaxillary gland androgen-regulated protein 3B | Unknown | 24 h, 48 h | Saliva |
| STATH | Statherin | General oral/dental health | 24 h, 48 h | Saliva |
| USP7 | Ubiquitin carboxyl-terminal hydrolase 7 | Protein homeostasis | Discharge | Neither |
| ZNF33A | Zinc finger protein 33A | Transcription factor | 48 h | Neither |

## Supplementary figures

**Fig S1.**
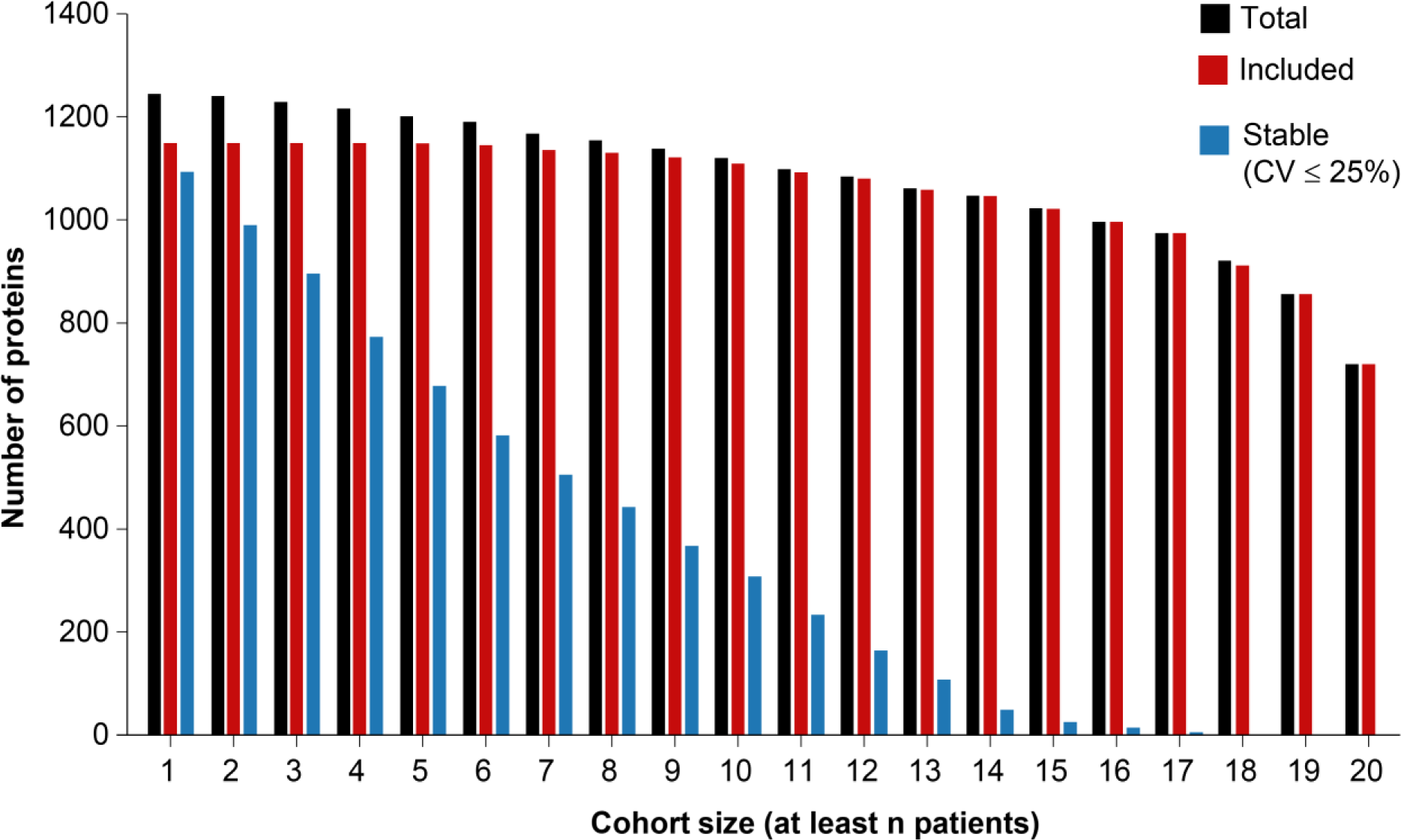
**Cumulative number of proteins detected in at least n participants**, filtered to 1% FDR. Tryptic peptides were analysed by C18-LC-DIA-MS/MS and analysed in FragPipe. Unique proteins were included in downstream analyses if detected in at least three time points within a participant and across at least three participants within the cohort. Proteins were defined as stable based on CV cut-off of 25% among uninfected samples.

**Fig S2.**
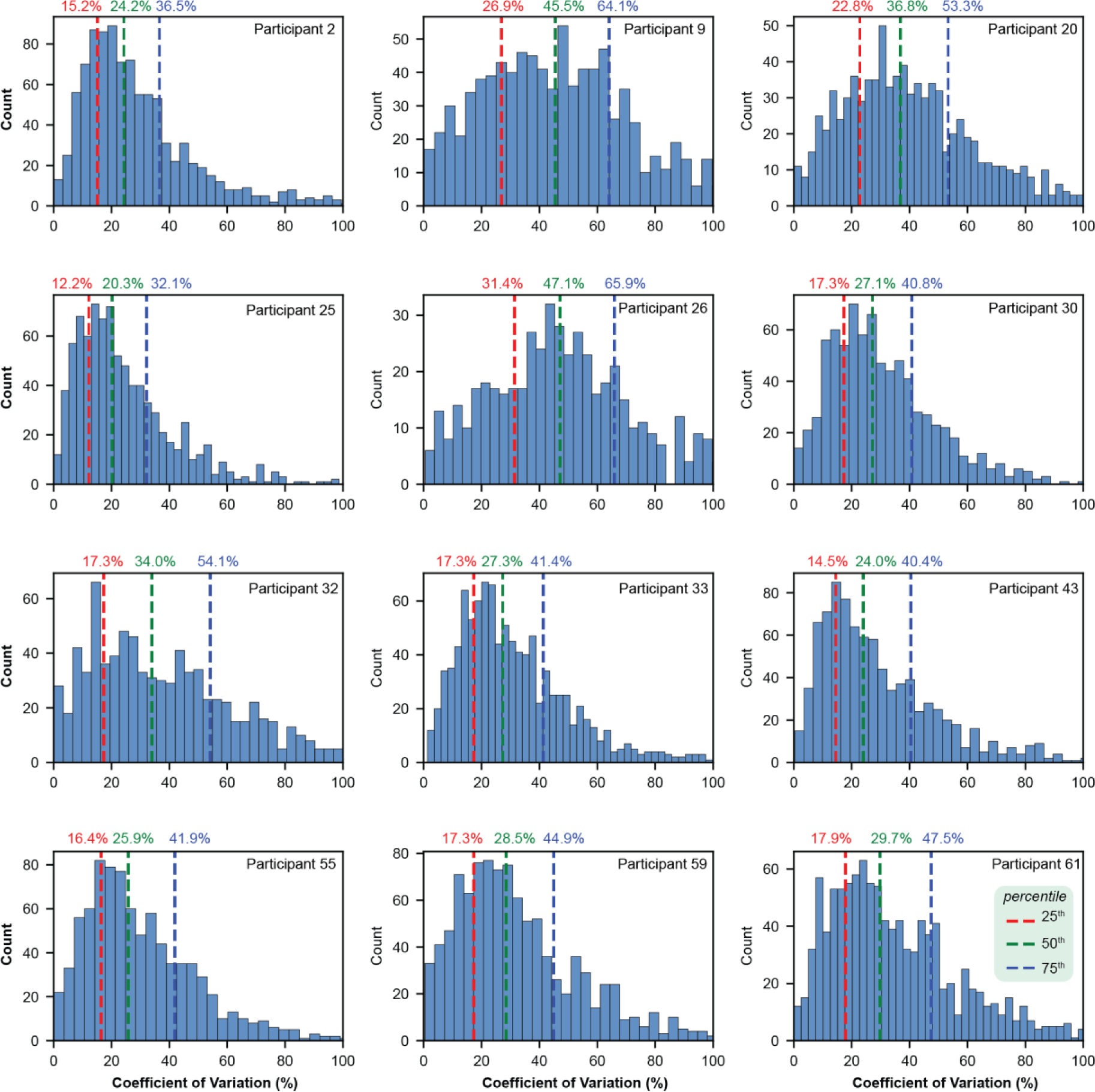
**Distribution of proteins by coefficient of variation** calculated across baseline (screening, evening before challege, 1 month post-discharge, 3 months post-discharge) time points, per individual. Twelve participants with sampling at all four uninfected time points are included.

**Fig S3.**
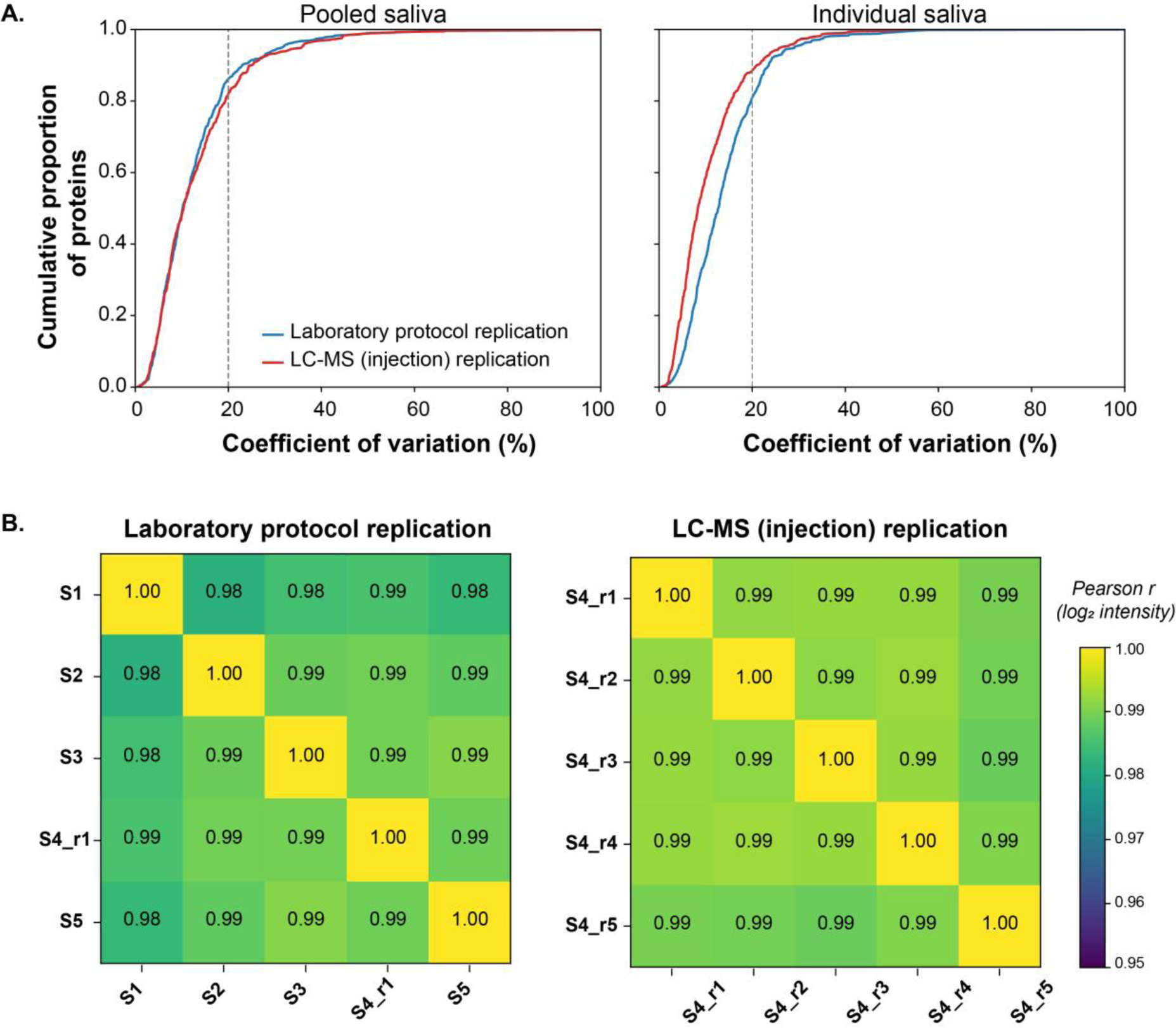
Assessment of technical reproducibility of saliva proteomics laboratory workflow and LC-MS analysis. (A) coefficient of variance on individual and pooled CHIVAS saliva, conducted according to established guidelines [66]. (B) Pearson correlation analysis of individual CHIVAS saliva. Pooled saliva comprised eight randomly selected CHIVAS samples. All modes of replication were performed with n=5 replicates.

**Fig S4.**
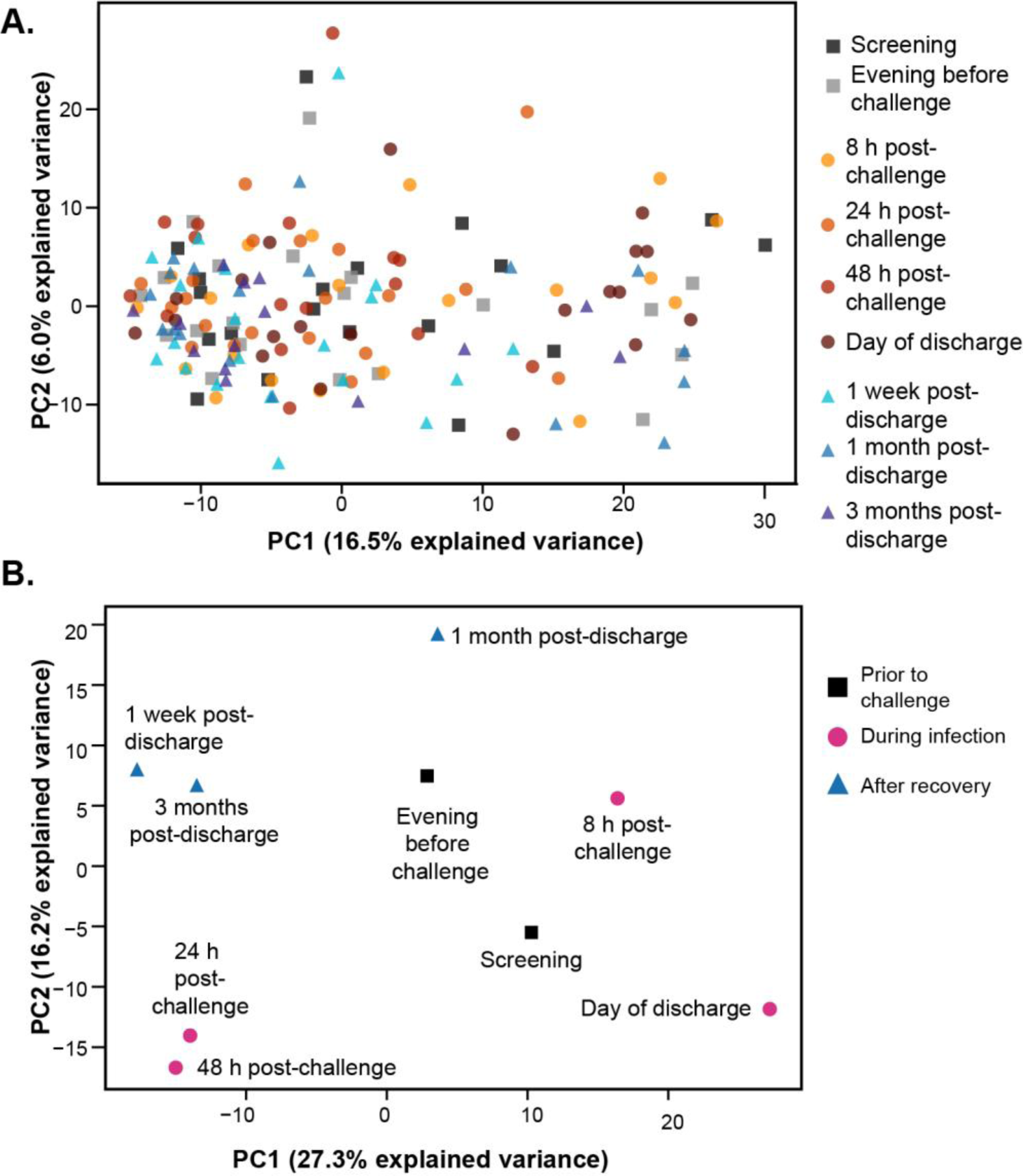
Principle component analysis of (A) individual and (B) time point-averaged saliva proteomes based on normalised DIANN abundance.

**Fig S5.**
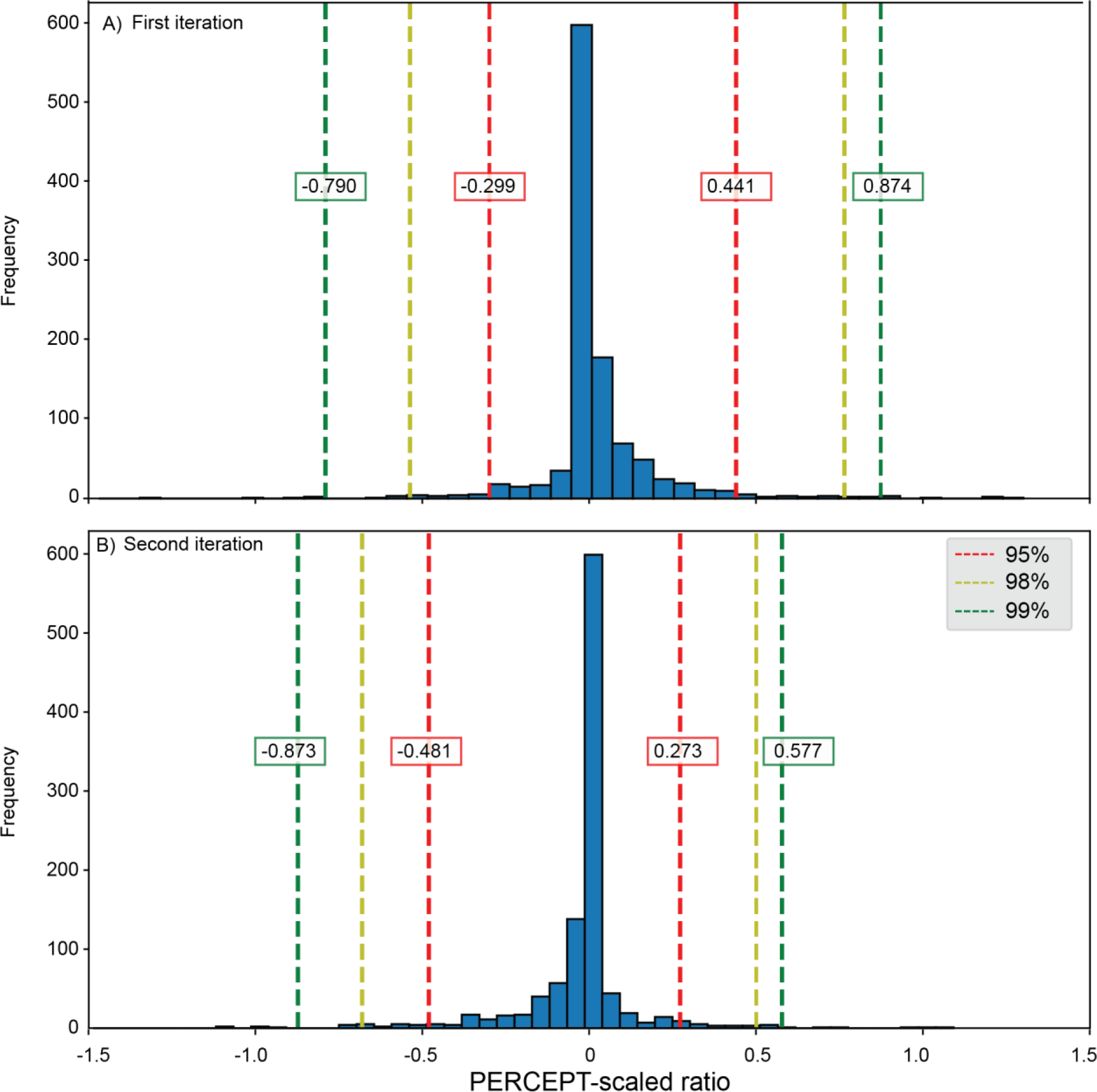
**Example PERCEPT-scaled ratios** calculated by pair-wise comparisons of salivary protein expression between randomly selected baseline (screening, evening before challege, 1 month post-discharge, 3 months post-discharge) time points. Two independent iterations are shown resulting in different threshold values, exhibiting both the variability of saliva and the necessity of repeated iteration. Thresholds were selected for downstream analysis by calculating the mean values of repeated iterations (n=100).

**Fig S6.**
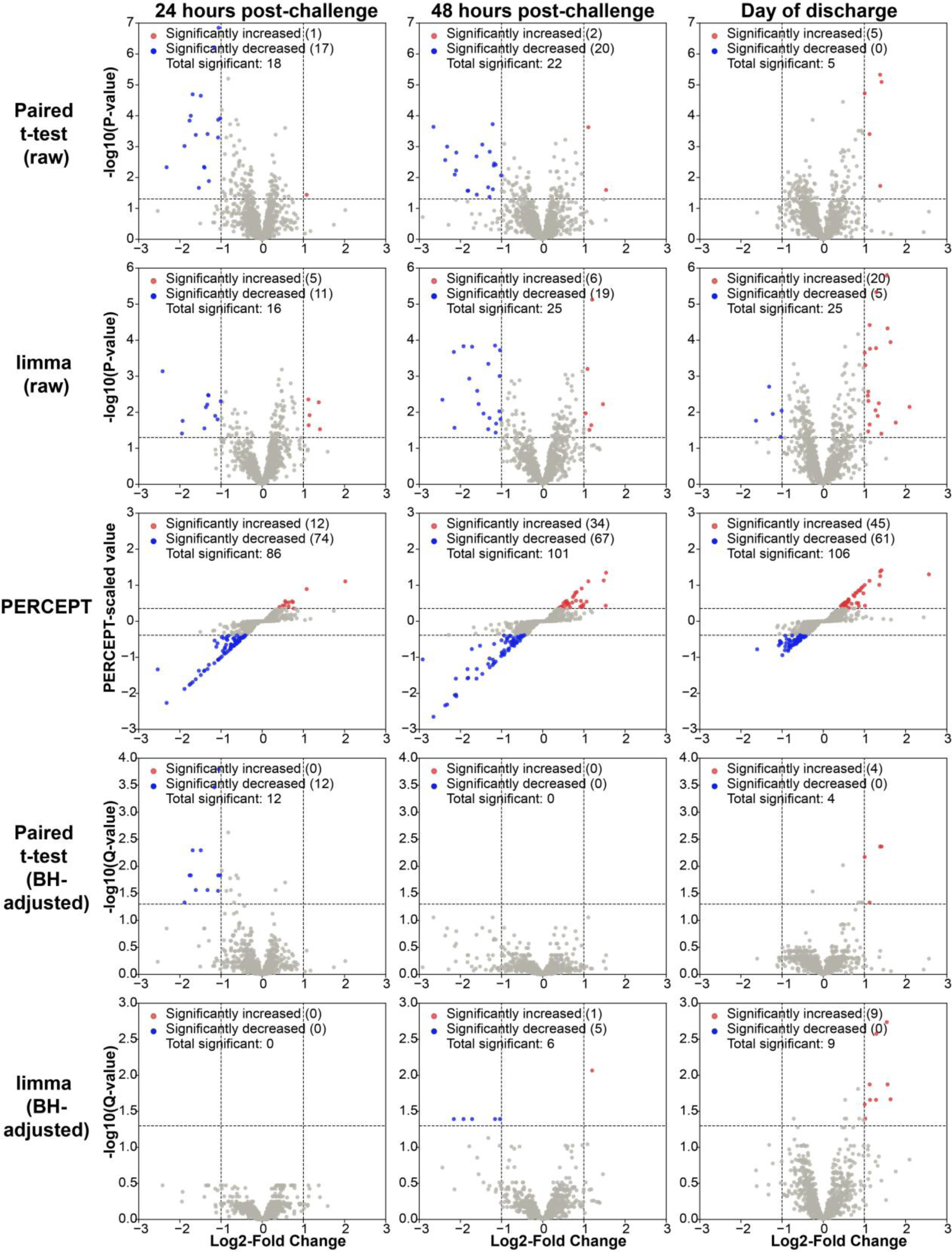
Concordance of five statistical approaches for differential abundance with and without Benjamini-Hochberg (BH) correction. A fixed threshold of |1| is used for t-test and limma.

**Fig S7.**
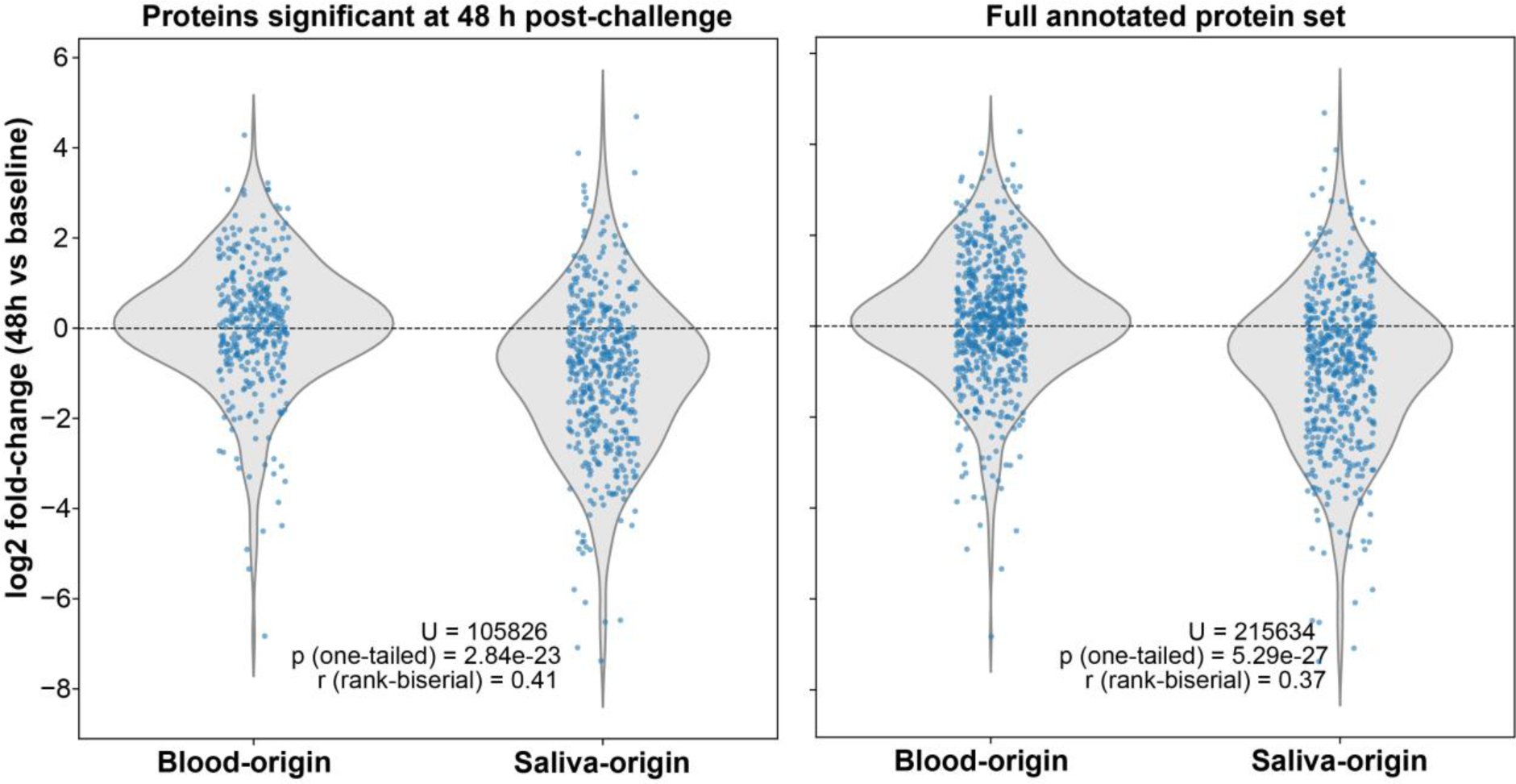
Comparison of trajectories of blood- and saliva-origin proteins. One-tailed Mann-Whitney U test was employed to test whether blood-origin proteins were overall increasing relative to saliva-origin proteins. Classification of proteins was achieved by detailed literature review.

**Figure S8.**
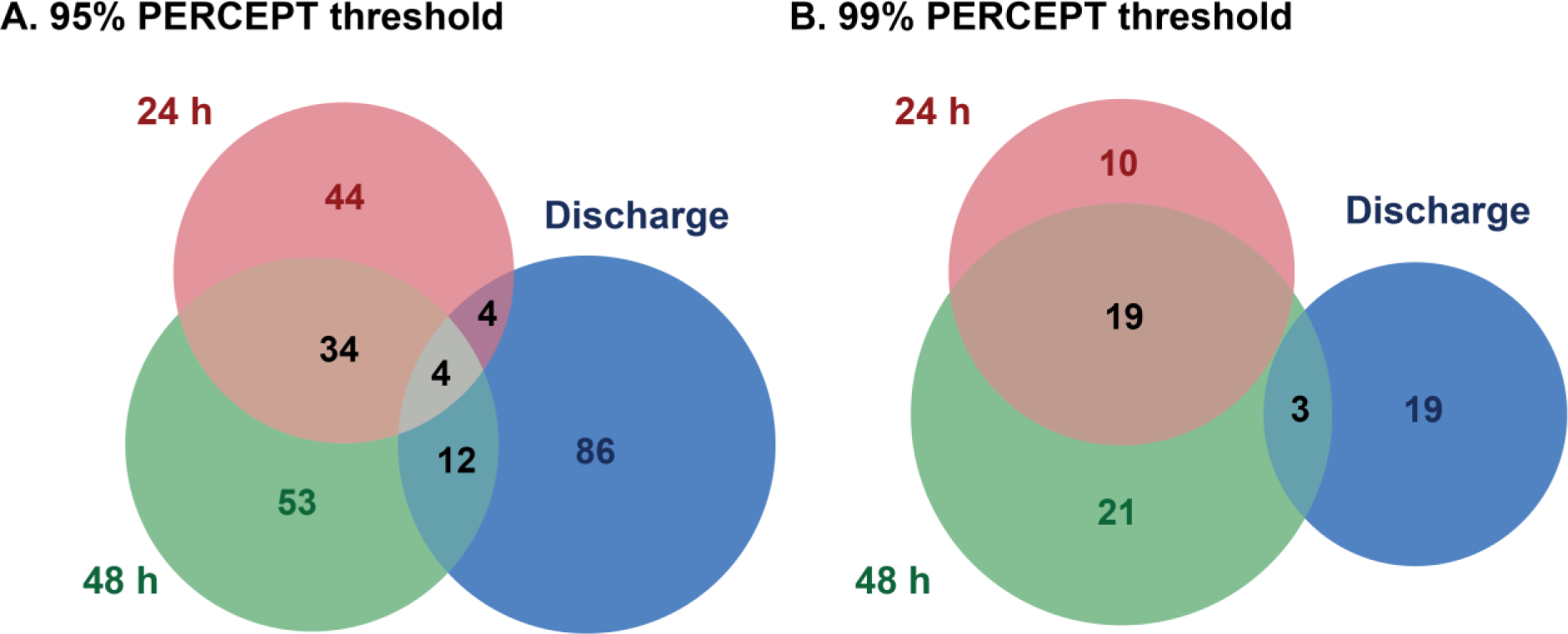
**Significant proteins in common between three post-infection time points** beyond (A) 95% of natural variance in PERCEPT-scaled values and (B) 99% of natural variance in PERCEPT-scaled values.

## References

[1] S. Brouwer, T. Rivera-Hernandez, B.F. Curren, N. Harbison-Price, D.M. De Oliveira, M.G. Jespersen, M.R. Davies, M.J. Walker, ‘Pathogenesis, epidemiology and control of Group A Streptococcus infection’, Nature Reviews Microbiology 21(7), (2023) 431–447.

[2] J.R. Carapetis, A.C. Steer, E.K. Mulholland, M. Weber, ‘The global burden of group A streptococcal diseases’, Lancet Infect. Dis. 5(11), (2005) 685–94.

[3] J. Vekemans, F. Gouvea-Reis, J.H. Kim, J.L. Excler, P.R. Smeesters, K.L. O’Brien, C.A. Van Beneden, A.C. Steer, J.R. Carapetis, D.C. Kaslow, ‘The Path to Group A Streptococcus Vaccines: World Health Organization Research and Development Technology Roadmap and Preferred Product Characteristics’, Clin. Infect. Dis. 69(5), (2019) 877–883.

[4] J. Osowicki, K.I. Azzopardi, C. Baker, C.S. Waddington, M. Pandey, T. Schuster, A. Grobler, A.C. Cheng, A.J. Pollard, J.S. McCarthy, M.F. Good, M.J. Walker, J.B. Dale, M.R. Batzloff, J.R. Carapetis, P.R. Smeesters, A.C. Steer, ‘Controlled human infection for vaccination against Streptococcus pyogenes (CHIVAS): Establishing a group A Streptococcus pharyngitis human infection study’, Vaccine 37(26), (2019) 3485–3494.

[5] J. Osowicki, K.I. Azzopardi, L. McIntyre, T. Rivera-Hernandez, C.Y. Ong, C. Baker, C.M. Gillen, M.J. Walker, P.R. Smeesters, M.R. Davies, A.C. Steer, ‘A Controlled Human Infection Model of Group A Streptococcus Pharyngitis: Which Strain and Why?’, mSphere 4(1), (2019).

[6] J. Osowicki, K.I. Azzopardi, L. Fabri, H.R. Frost, T. Rivera-Hernandez, M.R. Neeland, A.L. Whitcombe, A. Grobler, S.J. Gutman, C. Baker, J.M.F. Wong, J.D. Lickliter, C.S. Waddington, M. Pandey, T. Schuster, A.C. Cheng, A.J. Pollard, J.S. McCarthy, M.F. Good, J.B. Dale, M. Batzloff, N.J. Moreland, M.J. Walker, J.R. Carapetis, P.R. Smeesters, A.C. Steer, ‘A controlled human infection model of Streptococcus pyogenes pharyngitis (CHIVAS-M75): an observational, dose-finding study’, Lancet Microbe 2(7), (2021) e291–e299.

[7] L.V. Fabri, K.I. Azzopardi, J. Osowicki, H.R. Frost, P.R. Smeesters, A.C. Steer, ‘An emm-type specific qPCR to track bacterial load during experimental human Streptococcus pyogenes pharyngitis’, BMC Infect. Dis. 21(1), (2021) 463.

[8] J. Anderson, S. Imran, H.R. Frost, K.I. Azzopardi, S. Jalali, B. Novakovic, J. Osowicki, A.C. Steer, P.V. Licciardi, D.G. Pellicci, ‘Immune signature of acute pharyngitis in a Streptococcus pyogenes human challenge trial’, Nat Commun 13(1), (2022) 769.

[9] A.D. Indraratna, S. Mytton, A. Ricafrente, D. Millar, J. Gorman, K.I. Azzopardi, H.R. Frost, J. Osowicki, A.C. Steer, D. Skropeta, M.L. Sanderson-Smith, ‘A highly sensitive 3base assay for detecting Streptococcus pyogenes in saliva during controlled human pharyngitis’, Talanta 276 (2024) 126221.

[10] J. Osowicki, H.R. Frost, K.I. Azzopardi, A.L. Whitcombe, R. McGregor, L.H. Carlton, C. Baker, L. Fabri, M. Pandey, M.F. Good, J.R. Carapetis, M.J. Walker, P.R. Smeesters, P.V. Licciardi, N.J. Moreland, D.L. Hill, A.C. Steer, ‘Streptococcus pyogenes pharyngitis elicits diverse antibody responses to key vaccine antigens influenced by the imprint of past infections’, Nat Commun 15(1), (2024) 10506.

[11] S.L. Enkel, B. Wong, T.K. Hla, J. Pickering, T.C. Barnett, H.M.M. Thomas, N. Lansbury, J.R. Carapetis, J. Osowicki, A. Steer, L. Manning, A.C. Bowen, ‘Transmission potential of Streptococcus pyogenes during a controlled human infection trial of pharyngitis’, mSphere 9(10), (2024) e0051324.

[12] T.K. Hla, J. Osowicki, J.A. Marsh, S. Salman, M. Page-Sharp, O. Yoo, K. Azzopardi, M. Morici, K.T. Batty, R.K. Barr, S.L. Enkel, J. Kado, L. Hatchuel, A. Fulurija, J.S. McCarthy, T.L. Snelling, A.C. Steer, J. Carapetis, L. Manning, ‘Establishing the lowest penicillin concentration to prevent pharyngitis due to Streptococcus pyogenes using a human challenge model (CHIPS): a randomised, double-blind, placebo-controlled trial’, Lancet Microbe 6(5), (2025) 101038.

[13] H.A. Fryer, C. Pitt, H.R. Frost, N. Kandhari, S. Byars, P.S. Lim, R.L. Fettes, R.M. Lovett, T.T. Nguyen, K. Chheng, ‘Antibody responses against bacterial glycans affinity mature and diversify in germinal centers’, Immunity 59(3), (2026) 666–681. e5.

[14] A. Lepletier, D. Vedis, V. Ozberk, M. Anugraham, D. Bassette, A. Callcutt, H.R. Frost, K.I. Azzopardi, A.C. Steer, D. Kolarich, J. Osowicki, M.F. Good, M. Pandey, ‘Sub-clinical exposure to Streptococcus pyogenes drives the development of immunity’, bioRxiv (2025) 2025.07.02.662868.

[15] C.Z. Zhang, X.Q. Cheng, J.Y. Li, P. Zhang, P. Yi, X. Xu, X.D. Zhou, ‘Saliva in the diagnosis of diseases’, Int J Oral Sci 8(3), (2016) 133–7.

[16] S.J. Farnaud, O. Kosti, S.J. Getting, D. Renshaw, ‘Saliva: physiology and diagnostic potential in health and disease’, ScientificWorldJournal 10(1), (2010) 434–56.

[17] T. Nonaka, D.T.W. Wong, ‘Saliva Diagnostics’, Annu. Rev. Anal. Chem. (Palo Alto Calif.) 15(1), (2022) 107–121.

[18] M.A. Javaid, A.S. Ahmed, R. Durand, S.D. Tran, ‘Saliva as a diagnostic tool for oral and systemic diseases’, J Oral Biol Craniofac Res 6(1), (2016) 66–75.

[19] T.K. Fabian, P. Hermann, A. Beck, P. Fejerdy, G. Fabian, ‘Salivary defense proteins: their network and role in innate and acquired oral immunity’, Int. J. Mol. Sci. 13(4), (2012) 4295–4320.

[20] A.V. Amerongen, E.C. Veerman, ‘Saliva--the defender of the oral cavity’, Oral Dis. 8(1), (2002) 12–22.

[21] A.M.L. Pedersen, D. Belstrom, ‘The role of natural salivary defences in maintaining a healthy oral microbiota’, Journal of Dentistry 80 (2019) S3–S12.

[22] M. Boroumand, A. Olianas, T. Cabras, B. Manconi, D. Fanni, G. Faa, C. Desiderio, I. Messana, M. Castagnola, ‘Saliva, a bodily fluid with recognized and potential diagnostic applications’, J. Sep. Sci. 44(19), (2021) 3677–3690.

[23] A.V. Everest-Dass, D. Jin, M. Thaysen-Andersen, H. Nevalainen, D. Kolarich, N.H. Packer, ‘Comparative structural analysis of the glycosylation of salivary and buccal cell proteins: innate protection against infection by Candida albicans’, Glycobiology 22(11), (2012) 1465–79.

[24] A. Everest-Dass, S. Nersisyan, H. Maar, V. Novosad, J. Schroder-Schwarz, V. Freytag, J.L. Stuke, M.C. Beine, A. Schiecke, M.T. Haider, M. Kriegs, O. Elakad, H. Bohnenberger, L.C. Conradi, M. Raygorodskaya, L. Krause, M. von Itzstein, A. Tonevitsky, U. Schumacher, D. Maltseva, D. Wicklein, T. Lange, ‘Spontaneous metastasis xenograft models link CD44 isoform 4 to angiogenesis, hypoxia, EMT and mitochondria-related pathways in colorectal cancer’, Mol. Oncol. 18(1), (2024) 62–90.

[25] P. Sinitcyn, H. Hamzeiy, F. Salinas Soto, D. Itzhak, F. McCarthy, C. Wichmann, M. Steger, U. Ohmayer, U. Distler, S. Kaspar-Schoenefeld, N. Prianichnikov, S. Yilmaz, J.D. Rudolph, S. Tenzer, Y. Perez-Riverol, N. Nagaraj, S.J. Humphrey, J. Cox, ‘MaxDIA enables library-based and library-free data-independent acquisition proteomics’, Nat. Biotechnol. 39(12), (2021) 1563–1573.

[26] F. Yu, G.C. Teo, A.T. Kong, K. Frohlich, G.X. Li, V. Demichev, A.I. Nesvizhskii, ‘Analysis of DIA proteomics data using MSFragger-DIA and FragPipe computational platform’, Nat Commun 14(1), (2023) 4154.

[27] F.E. Dewhirst, T. Chen, J. Izard, B.J. Paster, A.C. Tanner, W.H. Yu, A. Lakshmanan, W.G. Wade, ‘The human oral microbiome’, J. Bacteriol. 192(19), (2010) 5002–17.

[28] A. Priyam, B.J. Woodcroft, V. Rai, I. Moghul, A. Munagala, F. Ter, H. Chowdhary, I. Pieniak, L.J. Maynard, M.A. Gibbins, H. Moon, A. Davis-Richardson, M. Uludag, N.S. Watson-Haigh, R. Challis, H. Nakamura, E. Favreau, E.A. Gomez, T. Pluskal, G. Leonard, W. Rumpf, Y. Wurm, ‘Sequenceserver: A Modern Graphical User Interface for Custom BLAST Databases’, Mol Biol Evol 36(12), (2019) 2922–2924.

[29] K. Hodge, S.T. Have, L. Hutton, A.I. Lamond, ‘Cleaning up the masses: exclusion lists to reduce contamination with HPLC-MS/MS’, J. Proteomics 88 (2013) 92–103.

[30] A.M. Frankenfield, J. Ni, M. Ahmed, L. Hao, ‘Protein Contaminants Matter: Building Universal Protein Contaminant Libraries for DDA and DIA Proteomics’, J. Proteome Res. 21(9), (2022) 2104–2113.

[31] D. Cox, D.M. Hatters, ‘PERCEPT: Replacing binary p-value thresholding with scaling for more nuanced identification of sample differences’, iScience 27(6), (2024) 109891.

[32] P. Shannon, A. Markiel, O. Ozier, N.S. Baliga, J.T. Wang, D. Ramage, N. Amin, B. Schwikowski, T. Ideker, ‘Cytoscape: a software environment for integrated models of biomolecular interaction networks’, Genome Res. 13(11), (2003) 2498–504.

[33] D. Szklarczyk, R. Kirsch, M. Koutrouli, K. Nastou, F. Mehryary, R. Hachilif, A.L. Gable, T. Fang, N.T. Doncheva, S. Pyysalo, P. Bork, L.J. Jensen, C. von Mering, ‘The STRING database in 2023: protein-protein association networks and functional enrichment analyses for any sequenced genome of interest’, Nucleic Acids Res. 51(D1), (2023) D638–D646.

[34] M.G. Ammari, C.R. Gresham, F.M. McCarthy, B. Nanduri, ‘HPIDB 2.0: a curated database for host-pathogen interactions’, Database (Oxford) 2016 (2016) baw103.

[35] D. Alonso-Lopez, F.J. Campos-Laborie, M.A. Gutierrez, L. Lambourne, M.A. Calderwood, M. Vidal, J. De Las Rivas, ‘APID database: redefining protein-protein interaction experimental evidences and binary interactomes’, Database (Oxford) 2019 (2019).

[36] S. Durmus Tekir, T. Cakir, E. Ardic, A.S. Sayilirbas, G. Konuk, M. Konuk, H. Sariyer, A. Ugurlu, I. Karadeniz, A. Ozgur, F.E. Sevilgen, K.O. Ulgen, ‘PHISTO: pathogen-host interaction search tool’, Bioinformatics 29(10), (2013) 1357–8.

[37] A. Calderone, L. Castagnoli, G. Cesareni, ‘mentha: a resource for browsing integrated protein-interaction networks’, Nat. Methods 10(8), (2013) 690–1.

[38] L. Happonen, S. Hauri, G. Svensson Birkedal, C. Karlsson, T. de Neergaard, H. Khakzad, P. Nordenfelt, M. Wikstrom, M. Wisniewska, L. Bjorck, L. Malmstrom, J. Malmstrom, ‘A quantitative Streptococcus pyogenes-human protein-protein interaction map reveals localization of opsonizing antibodies’, Nat Commun 10(1), (2019) 2727.

[39] M.E. Ritchie, B. Phipson, D. Wu, Y. Hu, C.W. Law, W. Shi, G.K. Smyth, ‘limma powers differential expression analyses for RNA-sequencing and microarray studies’, Nucleic Acids Res. 43(7), (2015) e47–e47.

[40] W.W. Lau, M. Hardt, Y.H. Zhang, M. Freire, S. Ruhl, ‘The Human Salivary Proteome Wiki: A Community-Driven Research Platform’, J. Dent. Res. 100(13), (2021) 1510–1519.

[41] M. Uhlen, L. Fagerberg, B.M. Hallstrom, C. Lindskog, P. Oksvold, A. Mardinoglu, A. Sivertsson, C. Kampf, E. Sjostedt, A. Asplund, I. Olsson, K. Edlund, E. Lundberg, S. Navani, C.A. Szigyarto, J. Odeberg, D. Djureinovic, J.O. Takanen, S. Hober, T. Alm, P.H. Edqvist, H. Berling, H. Tegel, J. Mulder, J. Rockberg, P. Nilsson, J.M. Schwenk, M. Hamsten, K. von Feilitzen, M. Forsberg, L. Persson, F. Johansson, M. Zwahlen, G. von Heijne, J. Nielsen, F. Ponten, ‘Proteomics. Tissue-based map of the human proteome’, Science 347(6220), (2015) 1260419.

[42] H. Elmongy, M. Abdel-Rehim, ‘Saliva as an alternative specimen to plasma for drug bioanalysis: A review’, TrAC, Trends Anal. Chem. 83 (2016) 70–79.

[43] J.A. Loo, W. Yan, P. Ramachandran, D.T. Wong, ‘Comparative human salivary and plasma proteomes’, J. Dent. Res. 89(10), (2010) 1016–23.

[44] C.S. Laxton, C. Peno, A.M. Hahn, O.M. Allicock, S. Perniciaro, A.L. Wyllie, ‘The potential of saliva as an accessible and sensitive sample type for the detection of respiratory pathogens and host immunity’, Lancet Microbe 4(10), (2023) e837–e850.

[45] D. Mehta, S. Scandola, R.G. Uhrig, ‘BoxCar and Library-Free Data-Independent Acquisition Substantially Improve the Depth, Range, and Completeness of Label-Free Quantitative Proteomics’, Anal. Chem. 94(2), (2022) 793–802.

[46] P.A. Wilmarth, M.A. Riviere, D.L. Rustvold, J.D. Lauten, T.E. Madden, L.L. David, ‘Two-dimensional liquid chromatography study of the human whole saliva proteome’, J. Proteome Res. 3(5), (2004) 1017–23.

[47] S. Hu, Y. Xie, P. Ramachandran, R.R. Ogorzalek Loo, Y. Li, J.A. Loo, D.T. Wong, ‘Large-scale identification of proteins in human salivary proteome by liquid chromatography/mass spectrometry and two-dimensional gel electrophoresis-mass spectrometry’, Proteomics 5(6), (2005) 1714–28.

[48] L.F.B. Miranda, C.V. Lima, R. Pagin, R.C. Costa, M.M.A. Pereira, E.D. de Avila, M. Bertolini, B. Retamal-Valdes, J.A. Shibli, M. Feres, V.A.R. Barao, J.G.S. Souza, ‘Effect of Processing Methods of Human Saliva on the Proteomic Profile and Protein-Mediated Biological Processes’, J. Proteome Res. 22(3), (2023) 857–870.

[49] L. Franco-Martinez, J.M. Gonzalez-Hernandez, A. Horvatic, N. Guillemin, J.J. Ceron, S. Martinez-Subiela, M.A. Sentandreu, M. Brkljacic, V. Mrljak, A. Tvarijonaviciute, P. Jimenez Reyes, ‘Differences on salivary proteome at rest and in response to an acute exercise in men and women: A pilot study’, J. Proteomics 214 (2020) 103629.

[50] M. Nissum, S. Kuhfuss, M. Hauptmann, C. Obermaier, U. Sukop, R. Wildgruber, G. Weber, C. Eckerskorn, J. Malmstrom, ‘Two-dimensional separation of human plasma proteins using iterative free-flow electrophoresis’, Proteomics 7(23), (2007) 4218–27.

[51] G. Saibaba, D. Rajesh, S. Muthukumar, G. Sathiyanarayanan, A.P. Aarthy, G. Archunan, ‘Salivary Proteome Profile of Women during Fertile Phase of Menstrual Cycle as Characterized by Mass Spectrometry’, Gynecol Minim Invasive Ther 10(4), (2021) 226-234.

[52] C.L. Chaiben, T.B.D. Batista, C.A.S. Penteado, M.C.M. Barbosa, T.M.O. Ventura, A. Dionizio, E.A.R. Rosa, M.A.R. Buzalaf, L.R. Azevedo-Alanis, ‘Salivary proteome analysis of crack cocaine dependents’, Arch. Oral Biol. 121 (2021) 104952.

[53] H. Xie, N.L. Rhodus, R.J. Griffin, J.V. Carlis, T.J. Griffin, ‘A Catalogue of Human Saliva Proteins Identified by Free Flow Electrophoresis-based Peptide Separation and Tandem Mass Spectrometry* S’, Mol. Cell. Proteomics 4(11), (2005) 1826–1830.

[54] K.C. Subbarao, G.S. Nattuthurai, S.K. Sundararajan, I. Sujith, J. Joseph, Y.P. Syedshah, ‘Gingival Crevicular Fluid: An Overview’, J. Pharm. Bioallied Sci. 11(Suppl 2), (2019) S135–S139.

[55] N.A. Ghallab, ‘Diagnostic potential and future directions of biomarkers in gingival crevicular fluid and saliva of periodontal diseases: Review of the current evidence’, Arch. Oral Biol. 87 (2018) 115–124.

[56] E. Gruys, M.J. Toussaint, T.A. Niewold, S.J. Koopmans, ‘Acute phase reaction and acute phase proteins’, J Zhejiang Univ Sci B 6(11), (2005) 1045–56.

[57] Y.Z. Szabo, D.C. Slavish, ‘Measuring salivary markers of inflammation in health research: A review of methodological considerations and best practices’, Psychoneuroendocrinology 124 (2021) 105069.

[58] C.N. LaRock, V. Nizet, ‘Inflammasome/IL-1beta Responses to Streptococcal Pathogens’, Front. Immunol. 6 (2015) 518.

[59] K. Wollein Waldetoft, T. Mohanty, C. Karlsson, M. Morgelin, I.M. Frick, J. Malmstrom, L. Bjorck, ‘Saliva-Induced Clotting Captures Streptococci: Novel Roles for Coagulation and Fibrinolysis in Host Defense and Immune Evasion’, Infect Immun 84(10), (2016) 2813–23.

[60] D.C. Nelson, J. Garbe, M. Collin, ‘Cysteine proteinase SpeB from Streptococcus pyogenes - a potent modifier of immunologically important host and bacterial proteins’, Biol Chem 392(12), (2011) 1077–88.

[61] H.S. Courtney, D.L. Hasty, J.B. Dale, ‘Anti-phagocytic mechanisms of Streptococcus pyogenes: binding of fibrinogen to M-related protein’, Mol. Microbiol. 59(3), (2006) 936–47.

[62] F. Carlsson, C. Sandin, G. Lindahl, ‘Human fibrinogen bound to Streptococcus pyogenes M protein inhibits complement deposition via the classical pathway’, Mol. Microbiol. 56(1), (2005) 28–39.

[63] O.D. Liang, K.T. Preissner, G.S. Chhatwal, ‘The hemopexin-type repeats of human vitronectin are recognized by Streptococcus pyogenes’, Biochem Biophys Res Commun 234(2), (1997) 445–9.

[64] B. Singh, Y.C. Su, K. Riesbeck, ‘Vitronectin in bacterial pathogenesis: a host protein used in complement escape and cellular invasion’, Mol. Microbiol. 78(3), (2010) 545–60.

[65] Y. Perez-Riverol, C. Bandla, Deepti J. Kundu, S. Kamatchinathan, J. Bai, S. Hewapathirana, Nithu S. John, A. Prakash, M. Walzer, S. Wang, Juan A. Vizcaíno, ‘The PRIDE database at 20 years: 2025 update’, Nucleic Acids Res. 53(D1), (2025) D543–D553.

[66] A.J. Brenes, ‘Calculating and Reporting Coefficients of Variation for DIA-Based Proteomics’, J. Proteome Res. 23(12), (2024) 5274–5278.

[67] M.C. Chambers, B. Maclean, R. Burke, D. Amodei, D.L. Ruderman, S. Neumann, L. Gatto, B. Fischer, B. Pratt, J. Egertson, K. Hoff, D. Kessner, N. Tasman, N. Shulman, B. Frewen, T.A. Baker, M.Y. Brusniak, C. Paulse, D. Creasy, L. Flashner, K. Kani, C. Moulding, S.L. Seymour, L.M. Nuwaysir, B. Lefebvre, F. Kuhlmann, J. Roark, P. Rainer, S. Detlev, T. Hemenway, A. Huhmer, J. Langridge, B. Connolly, T. Chadick, K. Holly, J. Eckels, E.W. Deutsch, R.L. Moritz, J.E. Katz, D.B. Agus, M. MacCoss, D.L. Tabb, P. Mallick, ‘A cross-platform toolkit for mass spectrometry and proteomics’, Nat. Biotechnol. 30(10), (2012) 918–20.

